# Multi-omic characterization of ILC and ILC-like cell lines as part of ILC cell line encyclopedia (ICLE) defines new models to study potential biomarkers and explore therapeutic opportunities

**DOI:** 10.1101/2023.09.26.559548

**Authors:** Osama Shiraz Shah, Fangyuan Chen, Abdalla Wedn, Anokhi Kashiparekh, Beth Knapick, Jian Chen, Laura Savariau, Ben Clifford, Jagmohan Hooda, Matthias Christgen, Jenny Xavier, Steffi Oesterreich, Adrian V. Lee

**Affiliations:** Integrative Systems Biology, School of Medicine, University of Pittsburgh; School of Medicine, Tsinghua University, Beijing, China; Department of Pharmacology & Chemical Biology, University of Pittsburgh; Department of Human Genetics, School of Public Health, University of Pittsburgh; Womens Cancer Research Center at UPMC Hillman Cancer Center and Magee Women’s Research Institute, Pittsburgh, PA; Department of Pharmacology & Chemical Biology, University of Pittsburgh, Pittsburgh, PA; Bionano Genomics, San Diego, California, USA; Institute of Pathology, Hannover Medical School, Hannover, Germany

## Abstract

Invasive lobular carcinoma (ILC), the most common histological “special type”, accounts for ∼10-15% of all BC diagnoses, is characterized by unique features such as E-cadherin loss/deficiency, lower grade, hormone receptor positivity, larger diffuse tumors, and specific metastatic patterns. Despite ILC being acknowledged as a disease with distinct biology that necessitates specialized and precision medicine treatments, the further exploration of its molecular alterations with the goal of discovering new treatments has been hindered due to the scarcity of well-characterized cell line models for studying this disease. To address this, we generated the ILC Cell Line Encyclopedia (ICLE), providing a comprehensive multi-omic characterization of ILC and ILC-like cell lines. Using consensus multi-omic subtyping, we confirmed luminal status of previously established ILC cell lines and uncovered additional ILC/ILC-like cell lines with luminal features for modeling ILC disease. Furthermore, most of these luminal ILC/ILC-like cell lines also showed RNA and copy number similarity to ILC patient tumors. Similarly, ILC/ILC-like cell lines also retained molecular alterations in key ILC genes at similar frequency to both primary and metastatic ILC tumors. Importantly, ILC/ILC-like cell lines recapitulated the *CDH1* alteration landscape of ILC patient tumors including enrichment of truncating mutations in and biallelic inactivation of *CDH1* gene. Using whole-genome optical mapping, we uncovered novel genomic-rearrangements including novel structural variations in *CDH1* and functional gene fusions and characterized breast cancer specific patterns of chromothripsis in chromosomes 8, 11 and 17. In addition, we systematically analyzed aberrant DNAm events and integrative analysis with RNA expression revealed epigenetic activation of *TFAP2B* – an emerging biomarker of lobular disease that is preferentially expressed in lobular disease. Finally, towards the goal of identifying novel druggable vulnerabilities in ILC, we analyzed publicly available RNAi loss of function breast cancer cell line datasets and revealed numerous putative vulnerabilities cytoskeletal components, focal adhesion and PI3K/AKT pathway in ILC/ILC-like vs NST cell lines.

In summary, we addressed the lack of suitable models to study E-cadherin deficient breast cancers by first collecting both established and putative ILC models, then characterizing them comprehensively to show their molecular similarity to patient tumors along with uncovering their novel multi-omic features as well as highlighting putative novel druggable vulnerabilities. Not only we expand the array of suitable E-cadherin deficient cell lines available for modelling human-ILC disease but also employ them for studying epigenetic activation of a putative lobular biomarker as well as identifying potential druggable vulnerabilities for this disease towards enabling precision medicine research for human-ILC.

## Introduction

Invasive lobular carcinoma (ILC), the most common histological “special type” of breast cancer (BC), accounts for approximately 10-15% of all BC diagnoses. It is characterized by unique features such as E-cadherin loss/deficiency, lower grade, hormone receptor positivity, larger diffuse tumors, distinctive metastatic patterns, and unique molecular alterations ^1–7^. Despite ILC being acknowledged as a disease with distinct biology that necessitates specialized and precision medicine treatments, the further exploration of its molecular alterations with the goal of discovering new treatments has been hindered due to the scarcity of well-characterized cell line models for studying this disease ^8–10^.

Pioneering large-scale cancer cell line characterization studies, including works by Ghandi et al ^11^, Marcotte et al ^12^, Iorio et al ^13^, and others ^14,15^, have revealed multi-omic molecular attributes and gene dependencies of diverse cancer cell lines. These investigations have facilitated a systematic selection of preclinical models that mirror the molecular characteristics of human cancers, enabling a more precise modeling of specific subtypes of breast cancer (such as the claudin-low and HER2-enriched types) and the exploration of their unique molecular alterations within the context of tumor initiation, progression, metastasis, resistance to treatments, and the development of new therapies ^12–14,16^. Comprehensive multiomic characterization of cancer cell lines has been invaluable in advancing precision medicine research ^14,16–23^. Similarly, precision medicine for ILC necessitates multi-omic characterization of ILC cell lines to guide rational selection of models to study this disease and elucidate the role of various molecular alterations in disease progression and treatment resistance, as well as to identify efficacious therapeutic strategies to guide clinical treatment decisions^8,9,24^.

However, a mere eight ILC cell lines have been established and made publicly available, of which only four (namely MDAMB134VI, MDAMB330, SUM44PE and BCK4) are established ER+ cell lines and are frequently utilized for modeling human-ILC disease ^8,9,24–27^. The remaining ILC cell lines have been inadequately characterized with only a few literature reports mentioning their use ^8,9^. Indeed, ILC cell lines, overall, lack comprehensive characterization, with limited scientific studies and scarce representation in large molecular characterization studies, such as the Cancer Cell Line Encyclopedia (CCLE) ^11^. Several factors can explain this limited availability and poor characterization; these include the lower incidence of ILC, the fewer number of ILC patients enrolled in clinical trials, the recent understanding of ILC as a unique disease, the fact that certain ILC cell lines are not yet publicly available/purchasable (including MA11 and WCRC25), and the intrinsic characteristics of ILC that make it challenging to culture them as cell lines ^2,8,9,28^. In pursuit of additional preclinical models to investigate human-ILC disease, a group of breast cancer cell lines (annotated as “carcinoma”, “adenocarcinoma” or “ductal carcinoma”) possessing genomic loss of E-cadherin or its binding partner α-catenin ‒ both events that impair E-cadherin function and result in disruption of adherens junctions – were proposed as potential models to study human-ILC. These were consequently labeled as “ILC-like” cell lines ^29^. However, since many of these ILC-like cell lines were established before ILC was recognized as a unique disease, it is still unclear whether these cell lines originated from human-ILC tumors or could serve as faithful models of this disease. In summary, both ILC and ILC-like cell lines lack comprehensive characterization and validation as faithful models for studying human-ILC disease, impeding the development of precision treatments for this disease.

To address this, we generated the ILC Cell Line Encyclopedia (ICLE), providing a comprehensive multi-omic characterization of ILC and ILC-like cell lines. ICLE cell lines exhibited pathognomonic E-cadherin deficiency and retained key ILC molecular alterations seen in patient tumors. Through whole-genome optical mapping, we characterized chromothripsis events and discovered novel genomic rearrangements, including new structural deletions in *CDH1* and functional gene fusions. Additionally, we systematically analyzed aberrant DNAm events and integrative analysis with RNA expression revealed the epigenetic regulation of *TFAP2B* – a putative lobular biomarker. Analysis of RNAi/CRISPR screens revealed druggable targets in PI3K/mTOR and focal adhesion pathways. Finally, we propose a qualitative and quantitative scheme to aid in rational selection of ILC/ILC-like cell lines for modeling human ILC disease. In summary, we generated ICLE – the first ILC cell line encyclopedia, highlighting key molecular features, cataloging druggable targets and presenting a scheme for rational-model-selection.

## Materials and Methods

### Multi-omic Profiling of ILC and ILC-like Cell Lines

#### ILC/ILC-like Cell Line Collection, Culturing and Extraction of RNA, DNA and protein lysates

All known *CDH1*/E-cadherin or *CTNNA1*/α-catenin deficient ILC/ILC-like cell lines reported in literature (ILC-like cell lines were reported in Michaut et al study ^29^) (Supplementary Table S1A). Two ILC-like cell lines (SKBR5 and ESVA-T) were not purchasable at the time of collection and therefore were excluded from the study. The final panel consisted of 17 ILC/ILC-like cell lines.

Most of these cells were purchased from vendors including CAMA1 (ATCC HTB-21), MDAMB134VI (ATCC HTB-23), MDAMB330 (ATCC HTB-127), SUM44PE (Asterand/BIOIVT HUMANSUM-0003016), UACC3133 (ATCC CRL-2988), HCC1187 (ATCC CRL-2322), HCC2218 (ATCC CRL-2343), MDAMB453 (ATCC HTB-131), MDAMB468 (ATCC HTB-132), SKBR3 (ATCC HTB-30), OCUBM (Riken RCB0881) and ZR7530 (ATCC CRL-1504). Several cell lines including 600MPE, HCC2185, IPH926, and BCK4 were kindly provided by Dr. Rachel Schiff (BCM) and Dr. Joe Gray (OHSU), Dr. John Minna (UTSW), Dr. Matthias Christgen (Hanover), and Dr. Britta Jacobson (CU ANSCHUTZ), respectively. WCRC25, a novel ILC cell line recently established in-house, now available through ABM (Catalog # T8074), was also included in the study ^30^. All cell lines were cultured using the recommended media conditions and underwent STR profiling based authentication at the Arizona genetics core. Please see Supplementary Table S1 for details on key publication, availability, media conditions of ICLE cell lines and other related information.

All cell lines were routinely assayed, after every few passages and before extraction of RNA, DNA, and protein lysates, to avoid Mycoplasma contamination using Lonza MycoAltert mycoplasma detection kit (Cat # LT07-118) and/or ABM mycoplasma PCR detection kit (Cat # G238) as per manufacturer’s instructions. Cell lines were grown in 150mm plates and used for RNA, DNA, and protein lysate collection when at ∼70% confluency. DNA extraction was performed using Qiagen QIAamp DNA mini kit (Cat # 56304) and RNA extraction using Qiagen RNeasy kit (Cat # 74004) as per vendor recommendations. All DNA/RNA samples showed high quality (260/280 ratio ∼1.8 for DNA and ∼2.0 for RNA) using NanoDrop (Cat # ND-2000). Protein lysates were prepared in accordance with the MD-Anderson RPPA core facility (https://www.mdanderson.org/research/research-resources/core-facilities/functional-proteomics-rppa-core/antibody-information-and-protocols.html) ^31^.

#### RNA sequencing (RNAseq) and data analysis

Extracted total RNA for all cell lines was sent to the UPMC Genome Center for RNAseq library preparation and sequencing. Briefly, RNA concentrations were quantified by ThermoFisher Qubit (Cat # Q33238) and their size distribution measured by Agilent TapeStation (Cat # G2991AA). Paired-end RNAseq libraries were prepared using KAPA poly-A RNA hyperPrep kit (Cat # 8098093702), and sequenced with NovaSeq 6000 for a target of 100 million reads/sample using S1 reagents for 300 cycles, 0.8 billion reads and total output of 250G (Cat # 20028317). Quality control of fastq files was performed using FastQC v0.11.9 ^32^ and summarized with multiQC ^33^. fastq files were aligned to hg38 reference transcriptome using STAR v2.7.5a ^34^ and gene count files generated using subread v2.0.1 featureCounts function ^35^. Gene counts were converted to counts per million (CPM) and log2 normalized using EdgeR ^36^ for downstream analyses in R ^37^. Ghandi et al 2019 (CCLE study) ^11^ RNAseq files were reprocessed using the pipeline described above and then gene counts merged with ICLE dataset to generate an integrated breast cancer cell line gene count matrix. Sensitivity to endocrine therapy was predicted using difference between ERpos and ERneg gene signatures as described previously ^38^.

#### Whole-exome sequencing (WES) and data analysis

Extracted DNA for all cancer cell lines (plus two control blood lines HCC1187BL and HCC2218BL) was sent to the UPMC Genome Center for WES library preparation and sequencing. Paired end libraries were prepared using Roche KAPA Hyper Plus Kit (Cat # 09075810001). Briefly, 500 ng of DNA was processed by fragmentation, enzymatic end repair, A-tailing and ligation. Quality control was done using Agilent Fragment analyzer. Exonic regions were captured using IDT xGen Exome Research Panel v1.0 (Cat # 1056114) with xGen Universal Blockers (Cat # 1075474) and Roche KAPA HyperCap workflow 3.0 (Roche) according to manufacturer protocol. The library size assessment was done using Fragment Analyzer (Agilent). Libraries with an average size of 400 bp (range: 200-600bp) were quantified by qPCR on the LightCycler 480 (Roche) using the KAPA qPCR quantification kit (Roche). The libraries were normalized and pooled as per manufacturer protocol (Illumina). Sequencing was performed using NovaSeq 6000 platform (Illumina) with 151 paired-end reads to an average target depth of 30-50X for germline and 125X coverage for somatic samples using S1 reagents for 300 cycles, 0.8 billion reads and total output of 250G (Cat # 20028317). Quality control of fastq files was performed using FastQC tool ^32^ and aligned to hg38 using BWA MEM tool ^39^. Variant calling was performed as per GATK best practices (Somatic short variant discovery (SNVs + Indels) – GATK (broadinstitute.org)). Briefly, GATK v4 MuTect2 pipeline ^40^ was used for calling single nucleotide variations (SNV) and indels with default parameters using HCC1187BL and HCC2218BL blood lines as normal control. All mutations were annotated using Ensemble variant effect predictor (VEP) ^41^ and converted to MAF format. Common polymorphisms (present in 1000genome database ^42^), silent mutations and mutations with allele frequency < 10% were removed before further analyses. Ghandi et al 2019 (CCLE study) MAF files filtered in similar way and merged with ICLE dataset to generate an integrated breast cancer cell line MAF file for downstream use. Tumor mutation burden (TMB) was computed using R package Maftools ^43^. Exonic deletions in *CDH1* and *CTNNA1* gene canonical isoforms were identified using samtools bedcov function.

#### Bionano optical genome mapping (OGM)

Frozen cell pellets containing at least 1.5 million cells were sent to Bionano Genomics Core, CA for OGM. Briefly, high molecular weight DNA (> 150 kbp) was extracted using Bionano SP blood and cell culture DNA isolation kit (Cat # 80042). DNA concentrations were measured using Qubit broad range assay (Cat # Q32850) and labelled using Bionano direct label and stain (DLS) labelling kit (Cat # 80005). Labelled DNA was transferred to a flow cell of a saphyr chip (Cat # 20366) and scanned repeatedly using a saphyr system with Bionano access server (Cat # 90067). The resulting images of molecules were converted to digital maps. These maps were aligned to reference genome maps to result in consensus whole-genome maps. Rare variant analysis and copy number analysis pipelines were used to detect structural variations, copy number variations and fusions at a minimum resolution of 5KB. The resulting structural variations were filtered against a panel of controls to remove pipeline artifacts and germline variations using Bionano access software. Filtered data was downloaded and further analyzed in R. All structural variations were categorized as one of the following: deletions (Del), insertions (Ins), inversions (Inv), duplications (Dup), inter-chromosomal translocations (Transloc-inter), and intra-chromosomal translocations (Transloc-intra). Correlation between structural variation count, mutation count and chromosome size were performed using Pearson correlation. Chromothripsis events were characterized using shatterseek tool using default parameters ^44^. Circos plots were generated using interacCricos ^45^.

#### Reverse Phase Protein Array (RPPA)

Protein lysates were sonicated and centrifuged at 14000rpm. Protein concentrations were determined by BCA. Then lysates were mixed with 4X SDS to have final concentration of 1ug/ml and then sent to RPPA core facility at MD Anderson Cancer Center, TX for RPPA. Briefly, the lysate was diluted and arrayed on nitrocellulose-coated slides and probed with antibodies to recognize signaling molecules. Signals are captured by DAB colorimetric reaction followed by spots are detection, background correction (using internal controls) and spot quality control. Curve fitting is used to quantify net relative concentrations and data is normalized for loading differences and other sources of variations. Final data comprised of ∼450 proteins and phospho-proteins. Marcotte et al 2016 (Neel Lab)^12^ normalized RPPA dataset was merged with normalized ICLE RPPA dataset to generate an integrated breast cancer cell RPPA dataset for downstream use.

#### Illumina DNA Methylation (DNAm) Array

Extracted DNA for all cell lines (except SKRB-3) was sent to University of Pittsburgh Genomic Research Core for DNA methylation profiling using Illumina Infinium Methylation EPIC BeadChip Kit (Cat # WG-317-1001). Briefly, DNA was quantified by Qubit and 500ng of DNA per sample was bisulfite converted using Zymo EZ DNA methylation kit (Cat # D5003) following vender recommendations. Bisulfite-converted DNA was hybridized to BeadChips and analyzed on an Illumina iScan as per vender recommendations.

Resulting raw IDAT files underwent quality control and preprocessing to result in B-values for 850K methylation probes using R package SeSAMe ^46^. B values were defined as SM/(SMLr+LrSU) for each methylation probe, where SM and SU represent signal intensities for methylated and unmethylated alleles. Probes with P value > 0.05 were masked as NA before downstream analysis. Probes were annotated for genomic context using Infinium annotations ^47^. Iorio et al 2016 (Sanger) ^13^ Infinium Methylation 450K array dataset was reprocessed using the pipeline described above and merged with ICLE EPIC array dataset to generate an integrated breast cancer DNAm dataset for downstream use.

#### Illumina CytoSNP Array

Extracted DNA for all cell lines (except SKRB-3) was sent to University of Pittsburgh Genomic Research Core for SNP profiling using Illumina Infinium CytoSNP-850K v1.2 BeadChip Kit (Cat # 20025644). Briefly, DNA was quantified using Qubit and 750 ng of DNA was amplified, fragmented, precipitated and hybridized to BeadChips and then analyzed on an Illumina iScan as per vendor protocol. Resulting raw IDAT files underwent quality control and preprocessing using Illumina Genome Studio version 2.0. Copy number (CN) analysis was performed using its CNVpartition v3.2.0 plugin. Copy number logR-ratio, B-allele frequencies (BAF) and genotype calls were exported and further analyzed in R. Copy number data was segmented using R package DNAcopy ^48^ and significantly amplified and deleted regions across samples were identified using GISTIC analysis ^49^. Marcotte et al 2016 (Neel Lab) ^12^ Illumina Omni Quad CN and SNP dataset was reprocessed using above pipeline and merged with ICLE CN and SNP dataset to generate integrated breast cancer cell line CN and SNP dataset. Fraction of genome altered (FGA) was computed using R package CINmeterics ^50^ by using segmented copy number.

The generated ICLE dataset data types are summarized in Supplementary Table S2, and the public datasets used for integration are summarized in Supplementary Table S3. The overlapping cell lines between ICLE and other public datasets showed strong genotyping similarity (Supplementary Figure S1). Mutations and copy number alterations were combined to define a single molecular/genomic alterations matrix for use in downstream analyses. All alterations were summarized into the following categories: mutation (MUT), mutation + LOH (MUT;LOH), mutation + single copy gain (MUT;GAIN), amplification (AMP) and deletions (DEL).

### Molecular Subtypes

PAM50 subtyping was performed using R package genefu ^51^. Briefly, this tool uses the method established in the Parker et al study ^52^. Briefly, Parker et al identified 50 genes that define transcriptional subtypes of breast cancer. Based on this training cohort of N=139, genefu computes PAM50 scores for a given input data. The PAM50 scores represent the correlation strength of each input sample to each PAM50 centroid in the training dataset. Final PAM50 subtype is defined as the one to which a given input sample has the strongest correlation score.

Consensus Clustering was performed individually on each integrated dataset to identify mRNA, RPPA and DNAm subtypes using ConsensusClusterPlus R ^53^. Briefly, ConsensusClusterPlus function was run on top 10% variable features with 1000 repeats and K = 9. Resulting clusters were evaluated based on their molecular features including E-cadherin, ER-A, PR, HER2, Cluadin-7 protein levels and ERpos/ERneg gene signature (adapted from Symmans et al ^38^) scores. ERpos/ERneg signature scores were defined as mean expression of CPM values of each gene in the signature. Clusters with similar molecular features were combined into a single cluster. Final subtypes were defined as following; Luminal (Lum) subtype comprised of cell lines with high ER-A/PR and low HER2/phospho-HER2 expression and had high ERpos gene signature score, HER2 subtype comprised of cell lines with low ER-A/PR and very high HER2/phospho-HER2 protein expression (due to HER2/ERBB2 gene amplifications) and had low ERpos gene signature score, Luminal/HER2 (Lum/HER2) subtype comprised of cell lines with high HER2/phospho-HER2 expression (due to HER2/ERBB2 gene amplifications) but unlike HER2 subtype showed similarity to Lum cluster and also had medium to high ERpos gene signature score, Basal subtype comprised of low ER-A/PR/HER2 cell lines and finally Claudin low (ClaudinLow) subtype comprised of basal cell lines which also had low levels of Claudin-7, a subset of ClaudinLow subtype cells showed down-regulation of E-cadherin and epithelial to mesenchymal (EMT) phenotype and were referred to as ClaudinLow/EMT. The results are shown in Supplementary Figure S2-S4 A-D and Figure 1B.

**Figure 1.**
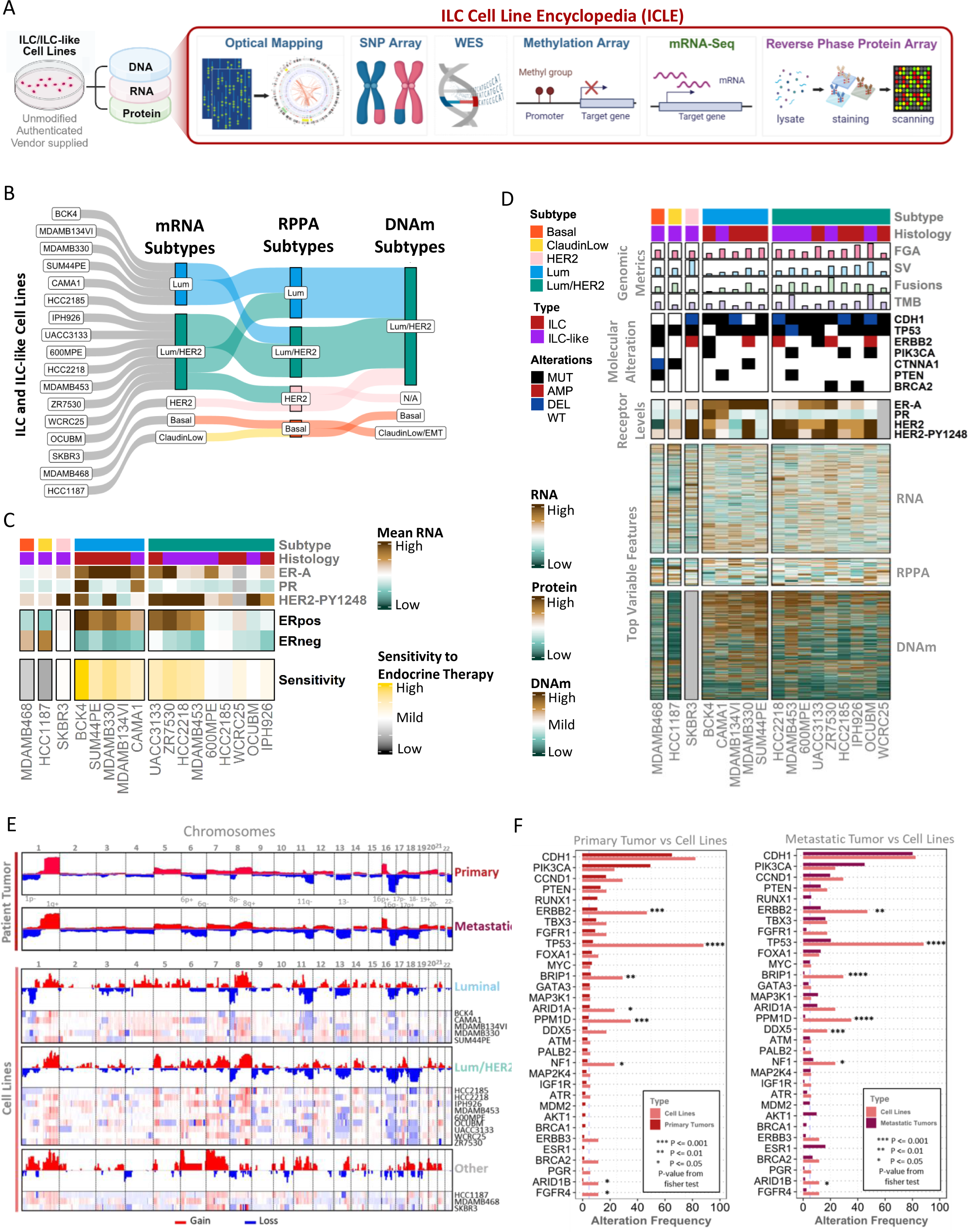
Multi-omic Characterization of ILC and ILC-like Cell Lines and Molecular Resemblance to Human-ILC Disease. (**A**) Overview of multi-omic profiling of 17 ILC and ILC-like cell lines using genomic (optical mapping, SNP array, whole-exome sequencing), epigenomic (DNA methylation array), transcriptomic (RNA sequencing) and proteomic (RPPA). (**B**) Multi-omic molecular subtypyes and their relationship with each other across each cell line. These subtypes were defined consensus clustering of mRNA, RPPA and DNA methylation (DNAm) datasets resulting in five mRNA subtypes (Luminal – Lum, Lum/HER2, Basal and CluadinLow), four RPPA subtypes (Lum, Lum/HER2, HER2, and Basal) and three DNAm subtypes (Lum/HER2, HER2, Basal, ClaudinLow/EMT and N/A for missing data on SKBR-3). (**C**) Hormone signaling activity and sensitivity to endocrine therapy (SET) in ILC/ILC-like cell lines. ERpos and ERneg gene signatures represent genes strongly associated with ER+ and ER-tumors, respectively. SET is defined as difference between mean of expression of ERpos and ERneg genes. (**D**) Overview of multi-omic features 17 cell lines clustered by mRNA subtypes. First panel highlights genomic metrics including fraction genome altered (FGA), structural variation (SV) count, fusion count and tumor mutation burden (TMB) and their values visualized as barplots. Second panel shows molecular alterations including two ILC driver alterations (CDH1 and CTNNA1) and other key ESCAT clinically actionable alterations. Third panel shows protein and levels of three receptors including ER-alpha (ER-A), PR, HER2 and phospho-protein levels of phosphorylated HER2-PY1248. Fourth, fifth and sixth panel show top variable multi-omic features including RNA (N = 5000), protein (N = 50) and DNAm (N = 5000). (**E**) Genome-wide linear plots showing copy number event frequencies across chromosomes in 1) N = 127 primary ILC tumors from TCGA, 2) N = 202 metastatic ILC tumors from MSK cohorts, 3) Luminal ILC/ILC-like cell lines, 4) Lum/HER2 ILC/ILC-like cell lines and 5) Other ILC/ILC-like cell lines. Copy gains are shown in red and copy loses in blue. Linear copy number plot of each cell line groups is followed by individual copy number plots of each cell line within that group. Key copy number alterations in patient tumors are listed below primary tumor copy number plot. (**F**) Barplot showing frequency of molecular alterations (mutations, amplifications and deletions) in various ILC associated genes and other clinically actionable breast cancer genes in primry ILC tumors vs ILC/ILC-like cell lines (left) and metastatic ILC tumors vs ILC/ILC-like cell lines (right). ILC associated genes include *CDH1, PTEN, TBX3* and *FOXA1* and others are clinically actionable breast cancer genes (defined by ESCAT – ESMO Scale for Clinical Actionability of molecular Targets). Fisher exact test was used to assess differences in frequency between comparison groups. All significant (p <= 0.05) differences are annotated while rest were non-significant.

### Molecular Resemblance to Patient Tumors & **CDH1** Alteration Landscape

The molecular resemblance of ILC/ILC-like cell lines to ILC patient tumors was assessed by comparing copy number profiles and molecular alteration frequency in key ILC genes. TCGA Primary ILC patient tumor ^3^ and MSK Metastatic ILC patient tumor ^54^ genome-wide segmented copy number, gene-level GISTIC values and mutation MAF datasets were acquired from cbioportal ^55^. Genome-wide segmented copy number profiles were visualized using integrative genomics viewer (IGV) ^56^ and key copy number events were visually compared between patient tumors and with cell lines. For a more quantitative comparison, we quantified similarity of tumor and cell line copy number profiles by Pearson correlation. Since there is a strong association between copy number values for genomically proximal loci the required data independence assumptions are invalidated. Instead, we used a sampling approach by randomly selecting 5 genes per genomic cytoband. Copy number values for these randomly selected genes in all cell lines were correlated with mean copy number profiles of TCGA ILC patient tumors (Figure 6B). Similarly, we quantified similarity of tumors and cell lines RNA expression using Pearson correlation. Since patient tumors also comprise of stroma, only the top 100 highly and lowly expressed genes in ILC tumors were used for correlation between all cell lines with mean expression of TCGA ILC patient tumors (Figure 6B).

Molecular/genomic alteration matrix for primary and metastatic ILC patient tumors was defined as described above for ILC/ILC-like cell lines. Frequency of molecular alterations (MUT, MUT;LOH, MUT;GAIN, AMP, and DEL) in ILC and other clinically actionable breast cancer genes defined by ESCAT (ESMO Scale for Clinical Actionability of Molecular Targets) ^6,57^ were compared between patient tumors and cell lines. Enrichment of specific gene molecular alterations in tumors vs ILC/ILC-like cell lines was computed using fisher exact test. Key oncogenic pathways used in Supplementary Figure S12 were adapted from a previous study ^58^.

To evaluate *CDH1* alteration landscape in patient tumors and cell lines, we performed integrative analysis multi-omic datasets including RNA expression, RPPA, mutations and copy number. TCGA ILC and NST patient tumors datasets were acquired from cbioportal ^3,55^ and integrated breast cell line datasets generated above were analyzed together. RNA and RPPA values were row normalized in patient and cell lines datasets separately. Only truncating (non-sense, nonstop, frame shift and splice site mutations) and missense mutations were considered in this analysis. *CDH1* mutation lollipop plot was generated using protein paint webtool (https://proteinpaint.stjude.org/) ^59^. Enrichment of *CDH1* biallelic alterations (MUT+LOH and Dual Loss) in ILC tumors and ILC/ILC cell lines vs NST tumors and NST cell lines was computed using fisher exact test.

### E-cadherin and p120-catenin Immunofluorescence

ILC/ILC-like cell line immunofluorescent analysis using E-cadherin and p120-catenin was performed as described in our previous study ^60^. Briefly, cells were plated at a density of 100,000–200,000 cells/well on glass coverslips in 24-well plates, fixed on ice in ice-cold methanol for 30 minutes and blocked in blocking buffer (0.3% Triton X-100, 5% BSA, 1X DPBS) for 1 hour at room temperature. Primary antibody incubation was performed overnight at 4°C: Cell Signaling Technology E-cadherin (Cat # 3195) and BD Biosciences p120 catenin (Cat # 610134). Secondary antibody incubation was done for 1 hour at room temperature followed by Thermo Fisher Scientific Hoechst 33342 (Cat # 62249) staining. Coverslips were mounted with Polysciences Aqua-Poly/Mount (Cat # 18606–20) and images were taken on a Nikon A1 confocal microscope. This experiment was performed by Dr. Laura Savariau.

### Aberrant DNA Methylation Events and Epigenetic Activation of**TFAP2B** Gene

To evaluate DNA methylation (DNAm) in breast cancer patient tumors, we utilized TCGA HM450K array dataset that was preprocessed using Sesame tool ^46^ by Silva et al in their recent publication ^61^. The DNAm instability (DMI) index per sample was computed by cumulative addition of DNAm deviations from control samples (tumor adjacent normal tissue). Briefly, DMI per sample was computed by adding all the differences between B-value of each probe compared that probes mean B-value in tumor adjacent normals. The resulting DMI scores were normalized against the tumor adjacent normals resulting in DMI z-scores.

To identify aberrant DNAm events in breast cancer patient tumors. Briefly, for each patient tumor the methylation intensity i.e., B-values of probe were compared to its distribution in tumor adjacent normal population. This allowed us to identify aberrantly methylated probes i.e., hyper- or hypo-methylated in tumor samples compared to normal samples. Hyper- and hypo-methylated probe frequency was compared between luminal A ILC vs NST patient tumors to identify aberrantly methylated probes in lobular disease (Figure 4C).

DNAm of all probes mapping to *TFAP2B* promoter region, defined as +/- 300bp of the transcription start site (TSS), and their effect on *TFAP2B* gene expression was investigated using Pearson correlation. Two probes mapping to *TFAP2B* gene TSS/exon 1 (cg25593948) and exon 2 (cg27260772) showed the strongest negative correlation with gene expression. Same probes were evaluated in TCGA HM450 dataset. Differential methylation analysis of TCGA Luminal-A ILC vs NST tumors using DNAm B-values was performed using R package Limma ^62^. Significant differentially methylated probes were defined as effect size > 0.1 and adjusted p-value < 0.05. These probes were mapped to genes using Infinium annotations ^47^. *TFAP2B* was the only gene with the highest number of differentially methylated probes (N = 12) as shown in Supplementary Figure S6B,C.

### Identification of Functional Fusions

Putative fusions identified from OGM were further evaluated to see whether they had any gain or loss of expression effects on one or both fused genes. This resulted in fusions that had functional effects on the fused genes, hereafter referred to as functional fusions.

Association of specific structural variation types (Del, Ins, Inv, Dup, Transloc-inter and Transloc- intra) and chromothripsis events with presence of functional fusions was computed using fisher exact test. Frequency of functional fusions in cell lines was compared those in patient tumor fusions using tumorfusion dataset (https://www.tumorfusions.org/) ^63^.

### Differential Gene Dependencies

To identify gene dependencies in ILC/ILC-like cell lines we utilized Marcotte el al 2016 (Neel Lab) RNAi loss of function screen dataset. There was data available for the following 10 ILC/ILC-like cell lines; MDAMB330, MDAMB134VI, CAMA1, HCC2218, OCUBM, SKBR3, MDAMB453, 600MPE, HCC2185 and SUM44PE and 12 luminal NST cell lines (Figure 19A). To perform differential dependency analysis, we utilized a consensus approach where differential dependencies were identified using three different algorithms. The original RNAi comprised of zGARP values. Same RNAi dataset reprocessed using DEMETER2 was acquired from DepMap (DepMap: The Cancer Dependency Map Project at Broad Institute ) ^16^. Both zGARP and DEMETER2 values underwent differential analysis using R package Limma ^62^. The final tool used for processing and differential analysis was R package siMEM ^12^ using default parameters. Consensus differential dependencies were curated from individual differential analyses (p-value < 0.05) using zGARP, DEMETER2 and siMEM. Consensus differential dependencies showing higher dependency in ILC/ILC-like cell lines were defined as high dependency and vice versa as low dependency genes. Geneset enrichment of high and low dependency genes using hypergeometric test was performed using R package hypeR ^64^. Druggable dependencies were identified by queries high and low dependency genes in drug gene interaction database dgidb.org/) ^65^.

## Data Availability

All data generated in this study is available in the supplementary files and on our local cbioportal instance (https://cbioportal.crc.pitt.edu/).

### Figures & Illustrations

Most bioinformatic analysis visualizations were generated using R packages ComplexHeatmap ^66^ and ggplot2 ^67^. Mutational lollipop plot was made using proteinpaint.stjude.org ^59^. These visualizations were aesthetically improved in power point to produce the final figures. Some illustrations including an overview of ICLE were prepared using BioRender.com.

## Results

### Multi-omic characterization of ILC and ILC-like Cell Lines

We initiated the generation of ILC cell line encyclopedia (ICLE) by collection of all available and acquirable *CDH1*/E-cadherin or *CTNNA1*/α-catenin deficient ILC and ILC-like cell lines. We successfully collected most cell lines except two ILC-like (SKBR5 and ESVA-T) which were not publicly purchasable at the time of project initiation. The final ICLE cell line panel included a total of 17 cell lines including 8 ILC and 9 ILC-like cell lines (Supplementary Table S1 ). ICLE cell lines underwent molecular characterization using various multi-omics assays including optical genome mapping (OGM), whole-exome sequencing (WES), RNA sequencing (mRNAseq), reverse-phase protein array (RPPA), DNA methylation (DNAm) array and SNP array (Figure 1A and Supplementary Table S1). ICLE datasets were integrated with public datasets containing NST cell lines for downstream analyses (Supplementary Table S2). Overlapping cell lines between ICLE and Marcotte et al (Neel Lab) datasets showed strong genotyping similarity suggesting same cell line variants were profiled in ours and previously published datasets (Supplementary Figure S1).

Consensus clustering mRNA, RPPA and DNAm datasets ( Supplementary Figure S2, S3 and S4) revealed concordant molecular subtypes across three datatypes with mRNA subtypes being the most granular and refined ( Figure 1B ). Using mRNA subtyping, five ILC/ILC-like cell lines were categorized as Luminal (Lum), 9 as luminal/HER2 (Lum/HER2) and the remaining 3 were grouped into HER2, Basal and Claudin Low (Claudin/Low) subtypes ( Figure 1B ). Both mRNA subtypes and PAM50 based intrinsic molecular subtypes successfully classified known ER+/luminal cell lines (such as MDAMB134VI, BCK4, SUM44PE, CAMA1 and MDAMB330) as Luminal (Supplementary Figure S2A,D). However, unlike mRNA subtyping, PAM50 based subtyping failed to characterize known SKBR3, a known HER2 cell line, correctly as HER2 subtype (Supplementary Figure S2A,D). Indeed, PAM50 based molecular subtyping was developed for patient tumor datasets ^51,52^ and may not be suitable for cell line datasets. Moreover, consensus clustering based mRNA subtypes provided more precise classification of a subset of cell lines (600MPE, UACC3133, HCC2185, WCRC25, IPH926, HCC2218, ZR7530, OCUBM, MDAMB453) with both luminal and HER2 features by grouping them into Lum/HER2 subtype. Since most ILC patient tumors are ER+ and luminal in nature, the best models would be ER+/luminal ILC/ILC-like cell line. Most ILC/ILC-like cell lines were Lum or Lum/HER2 subtype (Figure 1B) and notably, several Lum/HER2 ILC/ILC-like cell lines despite having low levels of ER-A/PR retained high expression of ER target genes associated with ER+ tumors (i.e., ERpos gene signature) as well as had high score for inferred sensitivity to endocrine therapy ( Figure 1C; Supplementary Figure S2A). Indeed, previous studies show that ILC cell lines derived from ER+ ILC tumors (such as IPH926) may lose ER expression but retain their luminal gene expression pattern ^68^.

**Figure 2.**
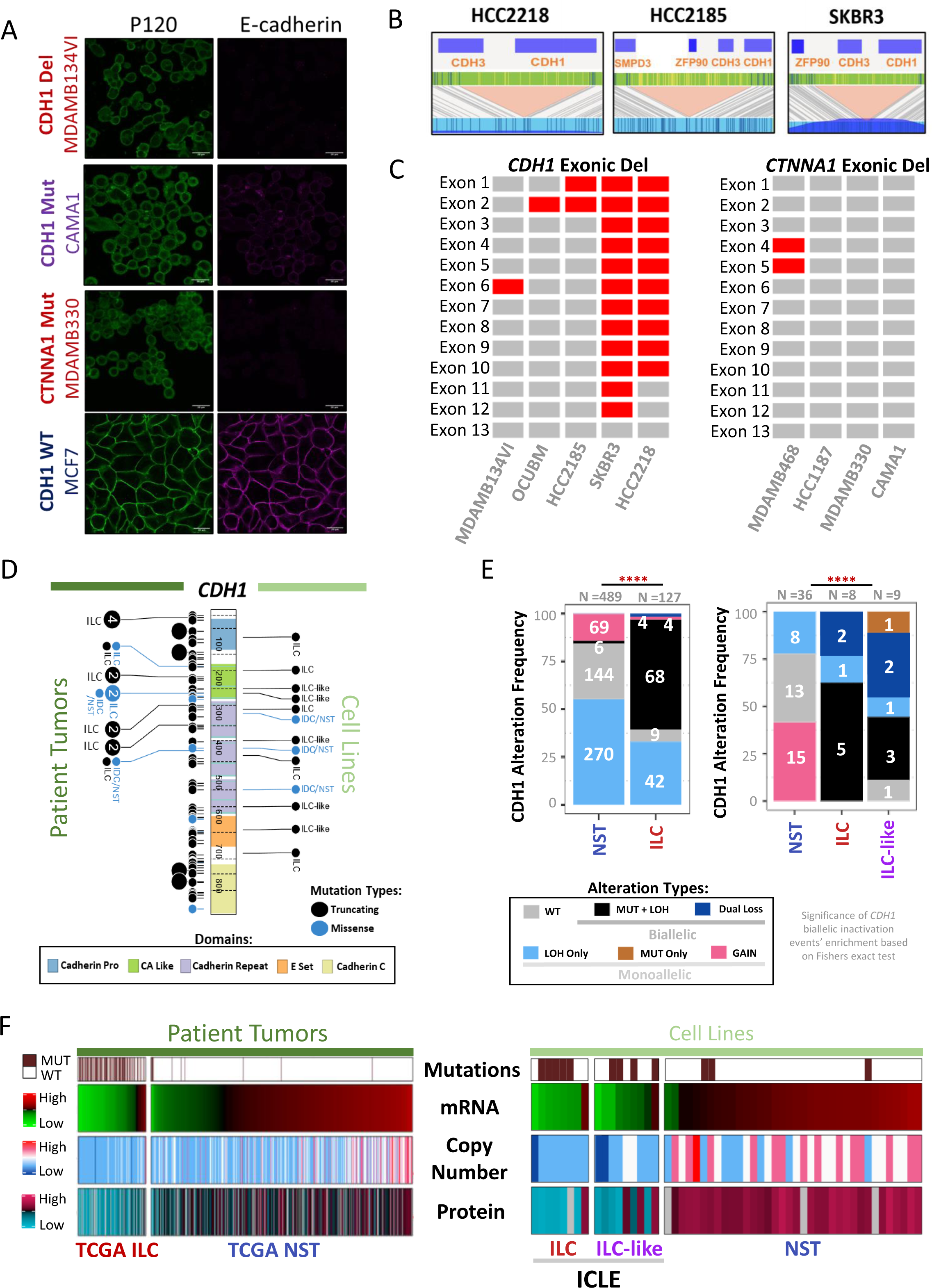
ILC and ILC-like Cell Lines Recapitulate CDH1 Alteration Landscape of Human-ILC Disease. (**A**) P120 and E-cadherin immunofluorescence of ILC/ILC-like cell lines including MDAMB134VI (ILC cell line harboring LOH + *CDH1* exonic deletion), CAMA1 (ILC-like cell line harboring *CDH1* mutation at 100% allele frequency) and MDAMB330 (a *CDH1* WT ILC cell line harboring *CTNNA1* mutation at 100% allele frequency) vs one representative *CDH1* WT NST cell lines (MCF7). (**B**) Novel structural variations resulting in *CDH1* deletions identified using optical genomic mapping in HCC2185, HCC2218 and SKBR3. (**C**) Exonic deletions identified in *CDH1* (E-cadherin) and *CTNNA1* (alpha-catenin) genes using WES dataset. Red tiles show exonic deletions. (**D**) Lollipop plot (generated using <u>proteinpaint.stjude.org</u>) showing truncating (black) and missense (blue) mutations across *CDH1* gene (including exons that form amino acid sequence and annotated with associated domains) in TCGA patient tumors (left) and ICLE/CCLE cell lines (right). The exon boundaries are shown by dotted lines, the numbers in each segment represent amino acid/protein length and the length of the lollipop represents the frequency of each mutation. (**E**) Barplots showing alteration frequency of *CDH1* gene in TCGA ILC vs NST patient tumors and ICLE ILC/ILC-like vs CCLE NST cell lines. Alterations are categorized as absence of alteration or wild-type (WT), mutation only (MUT Only), single allele copy number loss only (loss of heterozygosity – LOH Only), single allele copy number gain (GAIN), Mutation + LOH (MUT + LOH), and LOH plus exonic deletions or large structural deletions (Dual Loss). Association of *CDH1* biallelic inactivation in ILC vs NST tumors and cell lines was tested using fishers exact test (p-value < 0.0001, ****). (**F**) Heatmaps showing *CDH1* mutations, mRNA levels, copy number and protein levels in patient tumors (TCGA 127 ILC vs 489 NST) (left) and cell lines (ICLE 17 ILC and ILC-like vs CCLE 39 NST cell lines) (right).

**Figure 3.**
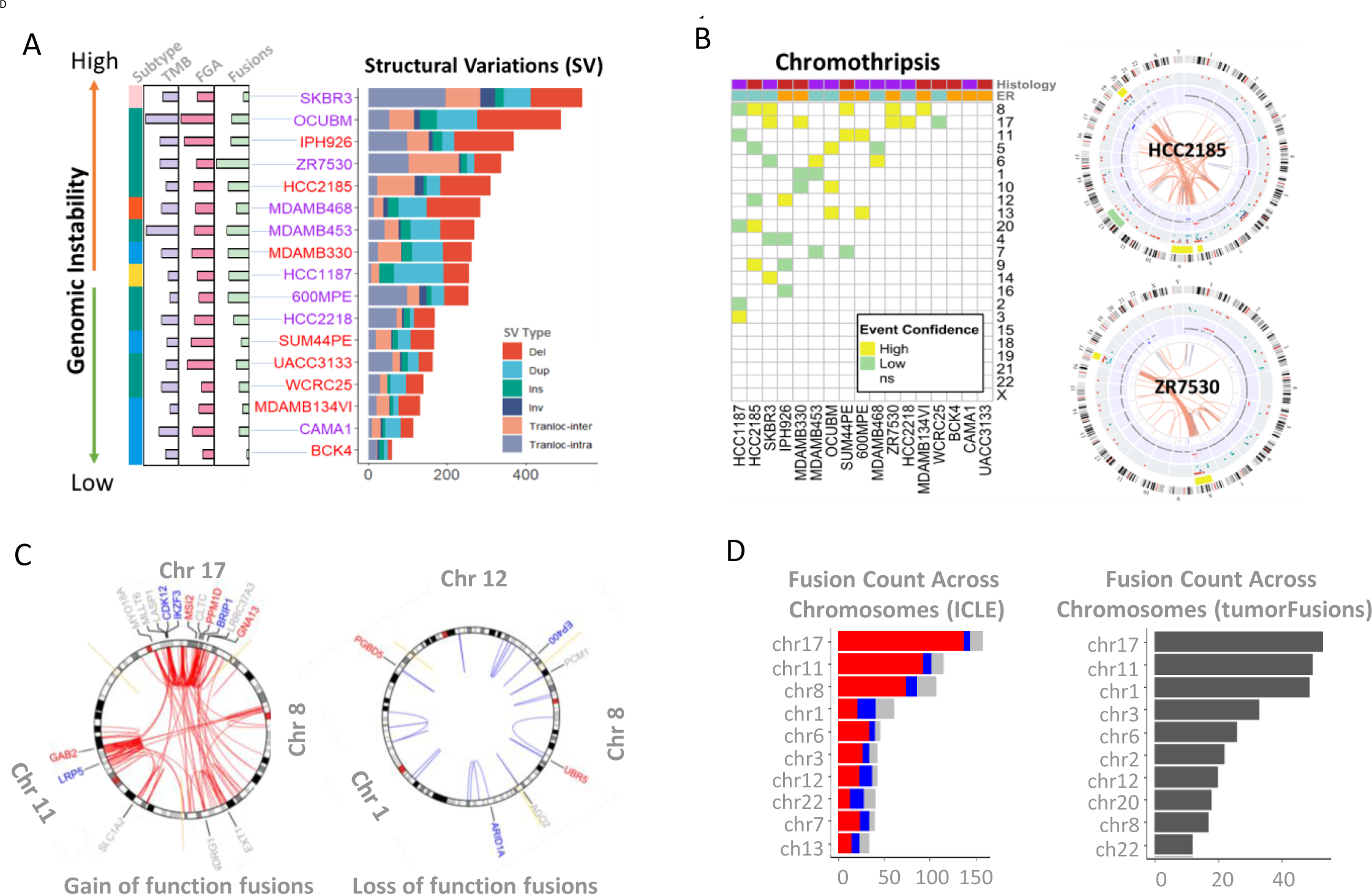
Optical Genome Mapping Unveils Structural Variations, Chromothripsis Events and Functional Gene Fusions. (**A**) Overview of structural variation types and their frequency across cell lines along with other metrics of genomic instability (TMB, FGA and # of fusions). SV types include deletions (del), duplications (dup), insertions (ins), inversions (inv), inter-chromosomal translocations (transloc-inter) and intra-chromosomal translocations (transloc-intra). (**B**) Heatmap (left) shows overview of chromosomal chromothripsis events predicted by shatterseek algorithm across cell lines. High and low confidence events are shown in yellow and green respectively. Circos plots (right) show SVs, CNVs and chromothripsis events in HCC2185 (ILC) and ZR750 (ILC-like) cell line. Details of each track are shown in supplementary figure 16. Briefly, track 1 hg19 cytoband, track 2 chromothripsis regions, track 3 SV track (excluding translocations), track 4 CNV track, track 5 gene annotations and track 6 translocations. (**C**) Shows circos plots highlighting translocation events in chromosome 8, 11 and 17 and chromosome 1, 8 and 12 resulting in gain of expression fusions (GEFs) and loss of expression fusions (LEFs), respectively. Only breast cancer driver genes involved in these gene fusions are shown. (**D**) Barplot showing frequency of fusion events across all chromosomes in ICLE (left) and Patient Tumors as reported in tumorfusions database (right). Chromosomes are ordered by decreasing order of fusion frequency with highest frequency of fusions in the chromosome 17 (left) and 15 (right).

**Figure 4.**
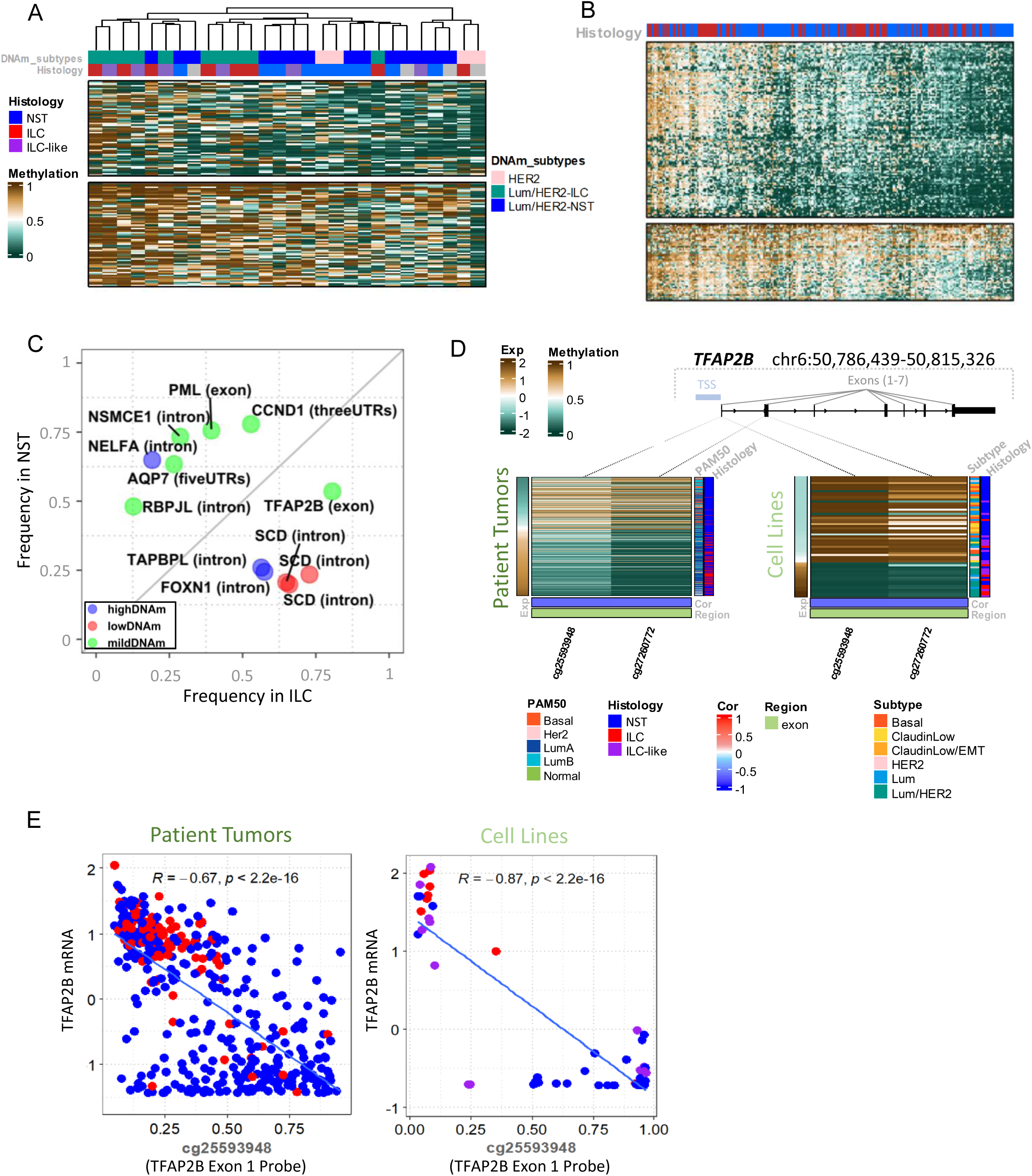
Aberrant DNA Methylation Events and Epigenetic Activation of TFAP2B. (**A**) Heatmap showing cell line – cell line similarity based on clustering using top 10% variable DNAm features across non-basal ILC/ILC-like and NST cell lines. (**B**) Heatmap showing sample – sample similarity based on clustering using top 10% variable DNAm features across TCGA luminal ILC and NST patient tumors. (**C**) Top aberrantly methylated probes and associated genes in TCGA luminal ILC vs NST patient tumors. Type of probe includes highDNAm (high DNAm in normal samples), midDNAm (mid DNAm in normal samples) and lowDNAm (low DNAm in normal samples). (**D**) Heatmaps showing regulation of *TFAP2B* gene expression via DNA methylation of probes mapping to its transcription start site (TSS)/Exon 1 (cg25593948) and Exon 2 (cg27260772) in both patient tumors (left) and cell lines (right). TFAP2B gene illustration shows its seven exons and location TSS. Patient tumors and cell lines are annotated by histology (ILC, NST and ILC-like) and intrinsic molecular subtypes (PAM50 for patient tumors and mRNA subtypes for cell lines). Probes are annotated by the region they map to and their Pearson correlation (cor) of DNA methylation levels with *TFAP2B* gene expression. (**E**) Scatter plots showing correlation between cg25593948 probe DNA methylation levels (B-value) with *TFAP2B* gene expression in patient tumors (left) and cell lines (right). The Pearson correlation coefficient (R) and p-value (p) are shown in each scatter plot.

Multi-omic features of profiled ILC/ILC-like cell lines are shown in Figure 1D and Supplementary Table S1 . All Lum cell lines showed high levels of ER-A and/or PR while low levels of HER2/phospho-HER2 (except MDMAB330, a HER2/ERBB2 amplified cell line). Similarly, Lum cell lines (except for MDAMB330) overall showed a lower frequency of structural variations (SV), fusions and tumor mutational burden (TMB). On the other hand, Lum/HER2 cell lines overall had low levels of ER-A/PR, but high levels of HER2/phosphor-HER2 (in cell lines with *HER2/ERBB2* mutations and amplifications). Moreover, they also had higher frequency of SVs, fusions and TMB with exception of WCRC25 which also showed lowest fraction of genome altered (FGA). Both OCUBM and MDAMB453 showed the highest TMB (>10 mutations per megabase) indicating a hypermutator phenotype. Non-luminal cell lines including SKBR3, HCC1187 and MDAMB468 lacked ER-A/PR levels and showed higher frequency of SVs and high TMB. ‘SKBR3 showed *ERBB2* amplification and consequently had higher levels HER2/phospho-HER2 while the other two cell lines lacked HER2 expression and were triple negative. All cell lines showed either *CDH1* or *CTNNA1* mutation as expected. Almost all showed *TP53* mutations or deletions except for BCK4 and ZR7530. Four cell lines showed alterations in *PIK3CA* and three in *PTEN* (both genes are components of oncogenic PI3K/AKT signaling) ^58^. Finally, only two cell lines had mutations in *BRCA2* including IPH926 (at 98% allele frequency indicating early event) and ZR7530 (at 32% allele frequency indicating later event in evolution).

### Molecular Resemblance to Human-ILC Disease

Although, only eight ILC/ILC-like cell lines had high ER-A/PR protein expression (Supplementary Table S1 ), many more showed high expression of ER target genes associated with ER+ tumors (i.e., ERpos gene signature) as well as had high score for inferred sensitivity to endocrine therapy (Figure 1C). Indeed, most (N = 14) ILC/ILC-like fell into luminal subtype in line with the observation that most ILC patient tumors are of luminal subtype^3^.

Breast cancers are driven by copy number alterations and major subtypes such as luminal, HER2 and basal subtypes each have distinct copy number profiles ^3,69,70^. Hence comparison of cell line copy number profiles to human tumors provides a valuable tool for assessment of genome-wide molecular resemblance ^12,71^. Towards this goal, we compared copy number profiles of Lum, Lum/HER2 and non-luminal/Other ILC/ILC-like cell lines with those of patient tumors. As most ILC/ILC-like cell lines were derived from metastatic lesion or ascites fluids ^8^, we included comparison with both primary and metastatic ILC patient tumors from TCGA ^3^ and MSK ^54^, respectively. Notably, both primary and metastatic ILC patient tumors showed very similar copy number profiles (Figure 1E) reminiscent of luminal tumors ^70^. Overall, Lum cell lines showed most similar copy number profiles to primary and metastatic ILC tumors followed by Lum/HER2 cell lines. Both Lum and Lum/HER2 subtype ILC/ILC-like cell lines showed high frequency of following chromosomal arm copy number alterations; 1p-, 1q+, 6p+, 8p+, 8q-, 11q-, 13p/q-, 16p+, 16q- (contains *CDH1* gene), 17q+ and 18p/q- events. Three non-luminal subtype cell lines were grouped together into “Other” subtype category. As expected, both MDAMB468 (basal subtype) and HCC1187 (claudinLow subtype) cell lines showed distinct copy number profiles and harbored 1p+ and 11p/q+ which were largely absent in other cell lines and patient tumors ^72^. On the other hand, SKBR3 (HER2 subtype) showed more resemblance to its luminal counterparts than its triple negative group members and harbored 1p-, 1q+,16q- and 18p/q-.

In addition to genome-wide copy number alterations, single nucleotide variations/alterations or mutations in particular genes also play a critical role in breast cancer initiation and progression. E.g., *CDH1* mutations are very common in ILC tumors and involved in tumorigenesis ^73,74^, while *GATA3* and *MAP3K1* mutations are more frequent in NST tumors ^3^ (Supplementary Figure S5A). In line with the “two hit” theory, tumor suppressor genes such as *CDH1* and *TP53* showed biallelic inactivation and harbored two molecular alterations e.g., truncating mutation in one allele and copy number loss of second allele or loss of heterozygosity (LOH) (Supplementary Figure S5A,B). To further assess molecular resemblance of ILC/ILC-like cell lines to human-ILC disease, we compared frequency of molecular alterations in key ILC genes and other clinically actionable genes between ILC/ILC-like cell lines vs primary or metastatic ILC tumors (Figure 1F). To identify whether frequency of key alterations in ILC/ILC-like cell lines is similar or distinct from patient tumors we performed fisher’s exact test.

Notably, the ILC/ILC-like cell lines retained molecular alterations, at similar frequency (fisher exact test non-significant) to primary and metastatic patient tumors, in key ILC genes including *CDH1* (82% in cell lines vs 65% in primary tumors and 80% in metastatic tumors), *PTEN* (18% in cell lines vs 13% in primary tumors and 13% in metastatic tumors), *TBX3* (18% in cell lines vs 16% in primary tumors and 10% in metastatic tumors) and *FOXA1* (12% in cell lines vs 7% in primary tumors and 13% in metastatic tumors). In addition, ILC/ILC-like cell lines also retained similar frequency (fisher exact test non-significant) of molecular alterations in numerous other key breast cancer and clinically actionable genes including *PIK3CA* (24% in cell lines vs 49% in primary tumors and 45% in metastatic tumors), *CCND1* (29% in cell lines vs 17% in primary tumors and 20% in metastatic tumors) and *FGFR1* (18% in cell lines vs 9% in primary tumors and 3% in metastatic tumors).

Unlike patient tumors, ILC/ILC-like cell lines showed higher frequency (and enrichment) of molecular alterations in following genes; *TP53* (88% in cell lines vs 8% in primary tumors and 20% in metastatic tumors), *ERBB2* (47% in cell lines vs 11% in primary tumors and 13% in metastatic tumors), *BRIP1* (29% in cell lines vs 6% in primary tumors and 1% in metastatic tumors), *PPM1D* (35% in cell lines vs 5% in primary tumors and 2% in metastatic tumors), *NF1* (24% in cell lines vs 4% in primary tumors and 7% in metastatic tumors), and *ARID1B* (12% in cell lines vs 1% in primary tumors and 2% in metastatic tumors). Interestingly, several of these genes lied on chromosome 17 (*TP53* on 17p13.1, *ERBB2* on 17q12, *BRIP1* on 17q23, and *PPM1D* on 17q23). Both *BRIP1* and *PPM1D* were the same 17q23.2 locus and were co-amplified in all altered cell lines (Supplementary Figure S5B).

In summary, ILC/ILC-like cell lines retained molecular alterations in key human ILC genes including *CDH1*, *PTEN*, *FOXA1* and *TBX3* and other breast cancer genes including *PIK3CA*, *CCND1* and *FGFR1* at similar frequency to ILC patient tumors. However, unlike ILC patient tumors, ILC/ILC-like cell lines showed higher frequency of alterations in several genes ( *TP53, ERBB2, BRIP1* and *PPM1D*) that are associated with genomic instability and DNA repair defects^3,69,70,72,75,76^.

All ILC/ILC-like cell lines showed mutations in *CDH1* (E-cadherin) or *CTNNA1* (α-catenin) genes (Figure 1D) and thus we hypothesized that these cell lines will have aberrant adherens junction like human-ILC disease. To confirm, we performed immunofluorescence staining analysis of 17 ILC/ILC-like cell lines (Supplementary Figure S7). As expected, ILC/ILC-like cell lines, similar to ILC patient tumors, showed aberrant adherens junctions defined by lack of membranous E-cadherin and/or cytoplasmic p120 staining ( Figure 2A ). Few ILC/ILC-like cell lines, including OCUBM, MDAMB134VII, HCC2218, SKBR3, HCC2185 and MDAMB468 did not harbor short mutations in the *CDH1* or *CTNNA1* gene and yet showed aberrant adherens junctions. We comprehensively analyzed all molecular alterations in the *CDH1* and *CTNNA1* gene loci to understand the molecular underpinnings of this phenotype in these cell lines. Using Bionano optical genome mapping, we found that three of these cell lines, HCC2185, HCC2218 and SKBR3, had novel large scale structural deletions in the *CDH1* gene along with LOH ( Figure 2B). While, both MDAMB13VII and OCUBM had exonic deletions in the *CDH1* genes along with LOH ( Figure 2C) and MDAMB468 had exonic deletion in the *CTNNA1* gene ( Figure 2C). These events explained the molecular underpinnings of aberrant adherens junctions phenotype in these seven cell lines.

Notably, almost all *CDH1* mutations occurring in ILC patient tumors and ILC/ILC-like cell lines were truncating in nature i.e., resulting in loss of E-cadherin protein, while those rarely seen in NST patient tumors and cell lines were miss-sense in nature and did not affect E-cadherin protein levels ( Figure 2D). Moreover, all *CDH1* mutations in ILC/ILC-like cell lines had near 100% allele frequency indicating these were clonal events that occurred early in evolution, while those in NST cell lines, except CAL-148 which had a clonal silent mutation, occurred at ∼50% allele frequency indicating sub-clonal events that occurred later in evolution (Supplementary Figure S5C). Overall, biallelic inactivation of *CDH1,* via either short mutation plus LOH (MUT+LOH) or large/exonic deletion plus LOH (Dual loss), was significantly enriched in both ILC vs NST patient tumors (p < 0.0001) and ILC/ILC-like vs NST cell lines (p < 0.0001) (Figure 2E). The multi-omic *CDH1* alterations landscape in breast cancer patient tumors and cell lines is illustrated shown in Figure 2F. Overall, ILC/ILC-like cell lines showed enrichment of both *CDH1* mutations, copy number loss, low mRNA and E-cadherin protein levels similar to like ILC patient tumors. Unlike ILC patient tumors and ILC/ILC-like cell lines, many NST patient tumors and cell lines showed *CDH1* copy number gains.

In summary, most ILC/ILC-like cell lines were of Lum or Lum/HER2 subtype and showed high activity of ERpos gene signature as well as high score for inferred sensitivity to endocrine therapy. Moreover, they showed similar copy number profiles to ILC patient tumors and retained molecular alterations in key ILC genes at similar frequency to patient tumors. Notably, most ILC/ILC-like cell lines recapitulated the *CDH1* alterations of human-ILC disease and showed concomitant disruption of adherens junctions.

### Characterization of Whole-Genome Structural Variations, Chromothripsis Events and Functional Gene Fusions

Whole-genome genomic rearrangements such as large structural variations and complex chromothripsis events have been largely uncharacterized in ILC/ILC-like cell lines. Towards achieving this goal, we performed optical genome mapping of ILC/ILC-like cell lines to identify structural variations, chromothripsis events and gene fusions (Figure 3A , Supplementary Figure S8).

Most frequent type of structural variations was deletions (Del, N = 1528), and least common was inversions (Inv, N = 166). However, if combined, intra-chromosomal and inter-chromosomal translocations together were the most abundant (N = 1942) type of structural variations. Overall, Lum ILC/ILC-like cell lines (except for MDAMB330) had lower frequency of structural variations than other subtypes with BCK4 having the fewest and SKBR3 (HER2 subtype) having the highest structural variations, indicating that Lum cell lines were genomically more stable than Lum/HER2 and non-luminal cell lines ( Figure 3A). Moreover, frequency of structural variations showed significant correlation (R = 0.58, p = 0.03) with frequency of mutations or TMB (Supplementary Figure S9A) suggesting that genomically unstable cell lines may have higher frequency of both structural variations and mutations. Moreover, unlike in the case of mutations, where its frequency was strongly correlated (R=0.65, p = 0.00081) with chromosomal size i.e., more mutations were found in longer chromosomes such as chromosome 1 (Supplementary Figure S9D,G), frequency of structural variations did not show significant correlation with chromosome size (R = 0.33, p = 0.13, Supplementary Figure S9C,F). However, there was a much stronger yet imperfect correlation between frequency of non-translocation structural variations and chromosomal size (R = 0.5, p = 0.014, Supplementary Figure S9E). This suggested that unlike mutations, structural rearrangements, especially translocations may selectively occur in specific genomic locations and may not just happen across the genome by chance.

Indeed, chromosomal translocation event breakpoints most frequently involved chromosome 8 (359 Transloc-intra, 392 Transloc-inter), 11 (120 Transloc-intra, 145 Transloc-inter) and 17 (161 Transloc-intra, 216 Transloc-inter) compared to all other chromosomes which on average had 20 Transloc-intra/chromosome and 39 Transloc-inter/chromosome (Supplementary Figure S8 and Supplementary Figure S9C). Moreover, many of these translocation events appeared clustered together and localized in nearby regions. This genomic instability was reminiscent of chromothripsis which is characterized by massive structural rearmaments occurring together and localized to isolated chromosomal regions that also oscillatory changes in copy number ^44^.

To comprehensively characterize these complex events, we employed shatterseek tool that identifies chromothripsis events by detecting intra-chromosomal translocations occurring in interleaved fashion and are accompanied by oscillatory copy number changes ^44^. We identified high and low confidence chromothripsis events across most cell lines except for BCK4, CAMA1 and UACC3133 ( Figure 3B and Supplementary Figure S8). High confidence chromothripsis events across cell lines were most frequent in chromosomes 8 (in 5 cell lines), 17 (4 cell lines) and 6/11/13 (2 cell lines each). Indeed, these findings are in line with previous reports showing that breast cancers have distinct chromothripsis patterns involving chromosomes 8, 11, and 17 ^77,78^.

Next, we identified N = 384 functional fusions from N = 525 putative fusions by selecting those that resulted in gain (N = 302) or loss (N = 82) of expression of its fusion partners (Supplementary Figure S10A). Like patient tumors, most functional fusions identified in ILC/ILC-like cell lines involved genes residing in chromosome 8 (N = 64 fusions), 11 (N = 63 fusions) and 17 (N = 89 fusions) (Figure 3D). Most of these functional fusions were gain of expression fusions (GEFs) which mostly resulted from structural rearrangements occurring in these three chromosomes (Figure 3C) and were strongly associated (fisher exact test, p = 0.0082) with inter-chromosomal translocation events. Moreover, GEFs also showed strong enrichment (fisher exact test, p = 0.029) in cell lines with chromosome 17 chromothripsis. On the other hand, loss of expression (LEFs) fusions involved genes residing on chromosome 1 (N = 10), 8 (N = 7) and 12 (N = 7) and were significantly associated with deletions events (fisher exact test, p = 0.0001).

Most common genes involved in fusions included *SHANK2* and *TENM4* on chromosome 11 and *BCAS3*, *GRB7*, *PPM1D* and *SKAP1* on chromosome 17 (Supplementary Figure S10B,C).

In summary, using optical genome mapping, we unveiled whole-genome structural variations landscape of ILC/ILC-like cell lines and characterized chromothripsis events and functional gene fusions.

### Aberrant DNA Methylation Events and Epigenetic Activation of**TFAP2B**

DNA methylation (DNAm) changes have been associated with breast cancer risk ^79–81^ and have been shown to be very different in distinct molecular subtypes of breast cancer ^82,83^. Moreover, identification of DNAm changes associated with distinct disease can reveal novel biomarkers for early detection and treatment response prediction ^84,85^. However, DNAm changes associated with lobular disease remain largely unexplored. Towards achieving this goal, we sought to explore DNAm in breast cancer lines and patient tumors towards identifying novel biomarkers of lobular disease.

Our initial exploration of DNAm profiles of breast cancer cell lines, as expected, revealed that cell lines cluster by their DNAm molecular subtypes (Supplementary Figure S4A,B). Notably, the DNAm subtypes closely resembled the mRNA subtypes (Supplementary Figure S4D). Next, we sought to quantify DNAm changes in cell lines vs. patient tumors by defining a DNAm instability index (DMI) which represents accumulative changes in DNAm across genome in cancer vs normal samples. We found that overall breast cancer cell lines showed more aberrant DNAm (high DMI) than patient tumors (Supplementary Figure S4E,F). Higher DMI in cell lines in part maybe driven by culture condition induced epigenetic changes^86,87^.

Next, we specifically looked at non-basal cell lines to better delineate the differences between NST and ILC histology. Clustering of non-basal cell lines by top variable DNAm B-values revealed that most ILC and ILC-like cell lines clustered separately from NST cell lines (Figure 4A). While clustering of luminal ILC and NST patient tumors by top variable DNAm B-value revealed that some ILC samples cluster away from NST samples while others cluster with them ( Figure 4B). To identify top aberrantly methylated regions in NST vs ILC patients, we took a systematic approach where we first identified non-variable methylated probes in normal samples. Then assessed whether these probes got aberrantly methylated (i.e., hyper- or hypo-methylated) in tumor samples compared to normal samples. Top aberrantly methylated probes and the associated genes in ILC vs NST patient tumors are shown in Figure 4C . Top aberrantly methylated probes and their associated genes in ILC included *TFAP2B, SCD, TAPBL* and *FOXN1*, while those in NST included *CCND1, PML, NSMCE1, NELFA* , *AQP7* and *RBPJL*. Notably, *TFAP2B* also came as the top hit in differential methylation analysis of in Luminal ILC vs NST patient tumors (Supplementary Figure S6B).

Importantly, *TFAP2B* gene encodes AP-2B, a novel mammary epithelial differentiation marker, which is preferentially expressed in ∼80% of ILC tumors ^88–90^. In vitro studies indicate that AP-2B controls tumor cell proliferation of ILC/ILC-like cell lines ^89^. AP-2B positivity is associated with overall favorable clinicopathological characteristics and prolonged relapse-free survival ^90,91^. Integrative analysis revealed a clear pattern of epigenetic regulation of *TFAP2B* gene expression by DNAm of the probes mapping to promoter/exon 1 and exon 2 of this gene (Figure 4D) in both breast cancer patient tumors and cell lines. *TFAP2B* epigenetic activation and its concomitant high expression were particularly observed in ILC patient tumors and ILC/ILC-like cell lines with exception of two ILC (MDAMB330 and SUM44PE) and four ILC-like (HCC2218, CAMA1, HCC1187 and MDAMB468) (Supplementary Figure S6A). Genomic alterations in *TFAP2B* were rare and were only seen in one ILC (amplification in UACC3133) and two NST (mutation in BT483 and CAL851) cell lines. Indeed, there was a strong negative Pearson correlation between *TFAP2B* gene expression and promoter/exon 1 probe methylation in breast cancer patient tumors and cell lines is shown in Figure 4E.

In summary, we explored the DNAm landscape of breast cancer patient tumors and cell lines and performed systematic and integrative analysis of breast cancer cell lines and patient tumor methylation datasets to reveal epigenetic activation of *TFAP2B*, an important emerging biomarker of lobular disease, in both ILC/ILC-like cell lines and ILC patient tumors.

### Cataloguing Gene Dependencies and Identifying Druggable Vulnerabilities

*CDH1* molecular alterations and E-cadherin deficiency is an attractive target for development of precision therapies for ILC disease. Several molecular studies have shown evidence that E-cadherin deficient cell lines have several druggable vulnerabilities including but not limited to GPCR signaling ^92,93^, PI3K/AKT and IGF1R pathway ^92,94^, and synthetic lethality with ROS1 inhibition ^93,95^. These studies have led to a currently ongoing precision medicine trial for ILC called ROSALINE which is investigating efficacy of ROS1 inhibitors in early setting of ER+ ILC disease ^96^. Despite these recent findings, a major bottleneck in comprehensive investigation of targeted therapies in ILC is lack of suitable preclinical models as described before. Indeed, most of the aforementioned studies and their findings are based on modeling E-cadherin defects in genetically modified cell lines and lack ILC context.

To overcome these limitations, we sought to identify genes that show higher dependency (i.e., essential for survival) in ILC/ILC-like cell lines vs NST cell lines. For this we employed publicly available RNAi loss of function screen datasets from Marcotte et al (Neel Lab) ^12^. To avoid contamination by subtype specific differences, we only used luminal cell lines for our analysis. We identified differential dependencies between 10 ILC/ILC-like vs 12 NST cell lines using RNAi datasets processed via three different algorithms to improve robustness of our findings (Figure 5A). Concordant differential dependencies across three algorithms were used to define consensus differential dependencies which included 98 high and 94 low dependency genes in ILC/ILC-like vs NST cell lines ( Figure 5B ). Over-representation analysis using KEGG database revealed that high dependency genes were associated with actin cytoskeleton, focal adhesion pathways and mTOR signaling, while low dependency genes were associated with cellular senescence and insulin signaling (Figure 5C). Both low dependency genes associated pathways were also identified in ingenuity pathway analysis (Supplementary Figure S11A,B). Select high dependency genes in the actin cytoskeleton, focal adhesion and PI3K/AKT/mTOR signaling are shown in Figure 5D. Both *PPP1R12B* and *ITGA11* genes were involved in more than one pathway. Next, we identified druggable gene dependencies by querying the identified gene dependencies (Figure 5B) in drug gene interaction database. This resulted in shortlisting of 17 high and 17 low dependency genes that were druggable and had targeted drugs against them (Figure 5E). Several of these druggable high dependency genes were components of actin cytoskeleton ( *CHRM4*), Focal adhesion ( *PDPK1*), PI3K/AKT/mTOR signaling ( *STK11*, *CCNE1*, *PDPK1*).

**Figure 5.**
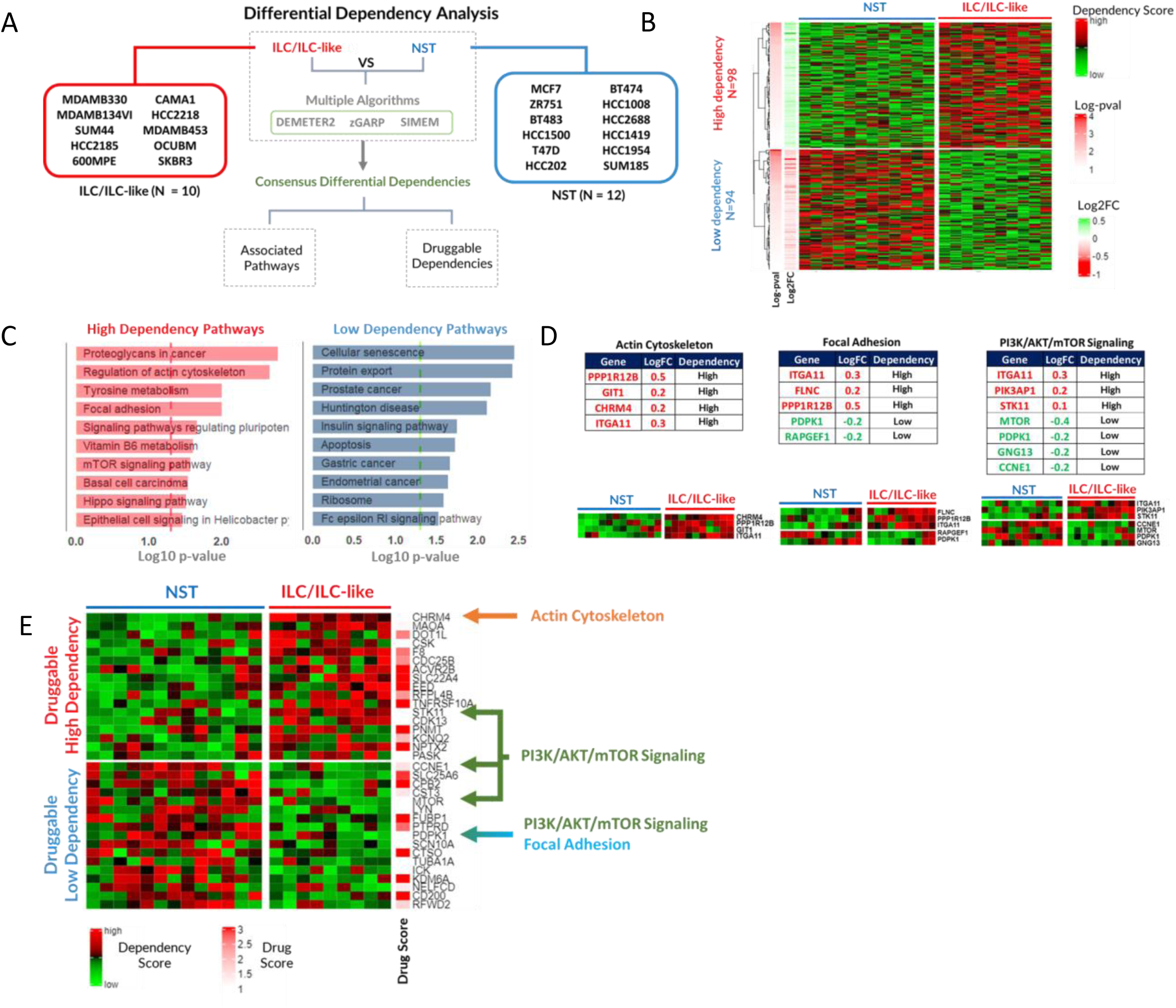
Identification of Differential Gene Dependencies in ILC/ILC-like vs NST Cell Lines. (**A**) Scheme of differential dependency analysis to identify ILC/ILC-like specific gene dependencies compared to NST cell lines. In total 10 ILC/ILC-like and 12 NST cell lines were selected with RNAi loss of function (LOF) screen data from Marcotte et al study (Neel Lab). Three differential analyses were performed using RNAi LOF screen dataset processed by three different algorithms (DEMETER2, zGARP and SIMEM). The overlapping findings from each differential analysis were used to define consensus differential dependencies. (**B**) Heatmap showing 98 high dependency genes and 94 low dependency genes identified from consensus differential dependencies in ILC/ILC-like vs NST cell lines identified. Differential dependency log2FC and log-pvalue from SIMEM algorithm are shown in left panels of the Heatmap. (**C**) Barplots showing KEGG pathways enriched in high (left) and low (right) dependency genesets identified in panel B. Enriched was computed using hypergeometric test. Barplot x-axis shows -log10 of p-value from hypergeometric test. (**D**) Key high dependency pathways and associated genes along with their log fold change (logFC) in the tables above and dependency scores in ILC/ILC-like vs NST cell lines in the heatmaps below. (**E**) Heatmap showing druggable high and low dependency genes identified by overlapping genes in panel B with druggable genes reported in drug gene interaction database (https://www.dgidb.org/) along with their reported druggability score (drug score). A higher score indicates strong literature evidence (# of reports) for a particular gene to be druggable/targetable by a specific drug. F) Overview of top druggable high dependency genes, putative drugs and drug scores.

**Figure 6.**
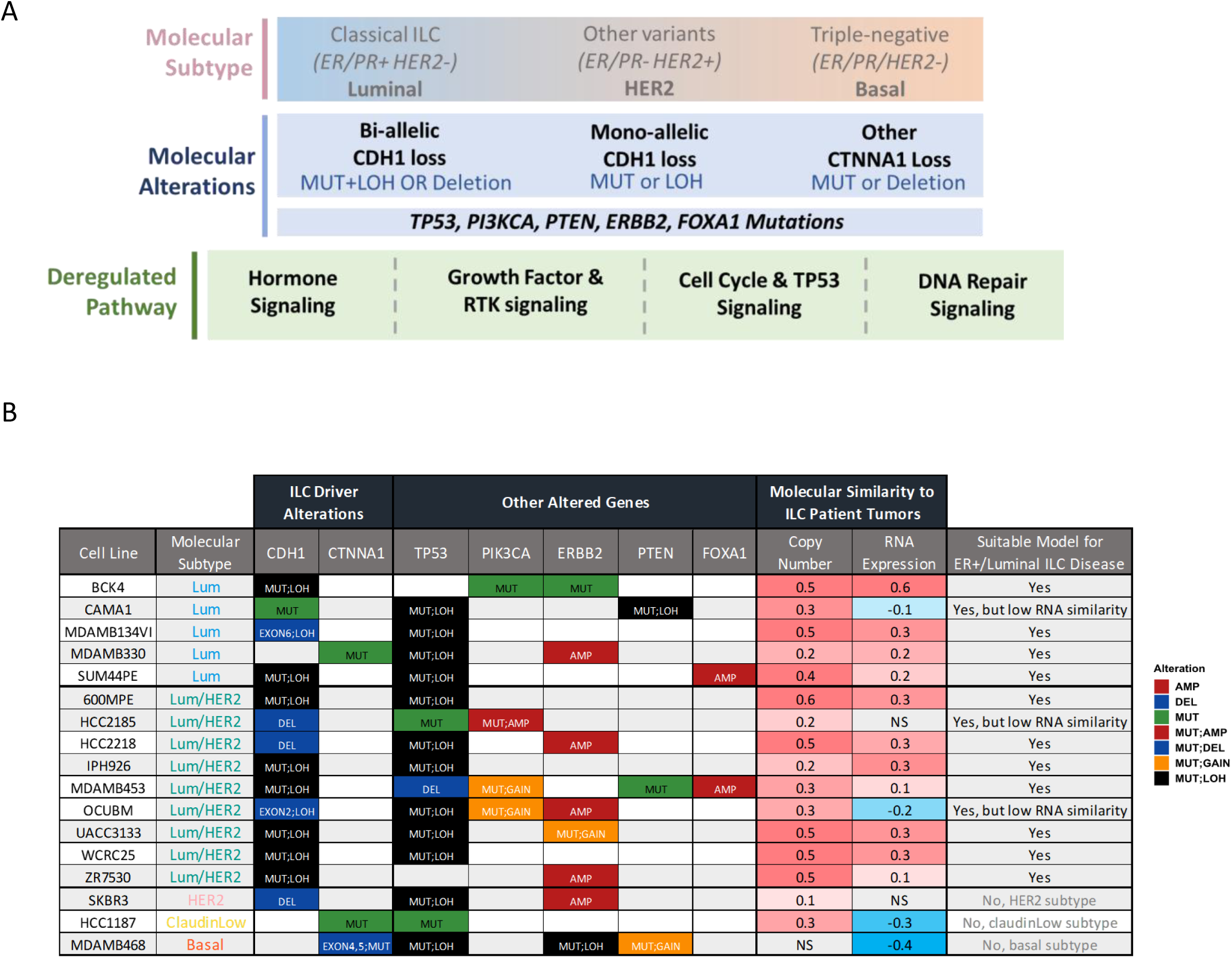
Scheme for Rational Selection of ILC/ILC-like Cell Lines for Modelling Human-ILC Disease.

In summary, our differential dependency analysis revealed numerous druggable vulnerabilities in ILC/ILC-like cell lines that hold potential to develop novel precision therapies for E-cadherin deficient breast cancers.

Scheme for Rational Selection of ILC/ILC-like Cell Lines for Modelling human-ILC disease ILC focused studies have majorly relied on a small subset of ILC/ILC-like cell lines for modeling this disease ( Supplementary Table S1 , see column of ILC focused PubMed reports). We sought to expand the list of suitable ILC/ILC-like cell lines available for modeling ER+/Luminal ILC disease by proposing a scheme for rational selection of these cell lines, summarizing their key multi-omic features considering this scheme and recommending suitable models.

The proposed scheme for rational selection of ILC/ILC-like cell lines and their key multi-omic features cell lines are summarized in Figure 6. This qualitative scheme is based on three criteria; 1) molecular subtype, 2) molecular alterations and 3) deregulated/altered pathways (Figure 6A). Breast cancer cell lines to model specific types of breast cancers are selected based on their molecular subtype. Since most ILC patient tumors are ER+ and luminal in nature, the best models would be ER+/luminal ILC/ILC-like cell line. Most ILC/ILC-like cell lines were Lum or Lum/HER2 subtype ( Figure 6B ) and notably, several Lum/HER2 ILC/ILC-like cell lines despite having low levels of ER-A/PR retained high expression of ER associated genes as shown by ERpos gene signature scores ( Figure 1C; Supplementary Figure S2A). Hence, overall, most luminal ILC/ILC-like cell lines would serve as useful models to study ER+/Luminal disease based on this criterion. On the other hand, non-luminal ILC/ILC-like cell lines such as HCC1187 (ClaudinLow), MDAMB468 (Basal) and SKBR3 (HER2) are not good models for ER+/luminal ILC disease but maybe useful for studying rare types of non-luminal ILC patients with similar molecular subtype or studying *CDH1* or *CTNNA1* defects in non-luminal breast cancers.

Next, it is important to select ILC/ILC-like cell lines based on their genetic context. Based on our initial selection, only ILC/ILC-like cell lines with alterations in *CDH1* or *CTNNA1* genes were selected. In addition to the ILC etiology defining alterations, several cell lines also had molecular alterations in less frequently altered ILC genes like *PTEN* (CAMA1, MDAMB453 and MDAMB468) and *FOXA1* (SUM44PE and MDAMB453). However, only 8% of ILC patient tumors have *TP53* molecular alterations, these very present in almost all ILC/ILC-like cell lines except BCK4 and ZR7530, making them suitable models to study *TP53* WT ILC tumors. Conversely, most (49%) of the ILC patient tumors have *PIK3CA* molecular alterations, but these were less frequent in ILC/ILC-like cell lines (which included BCK4, MDAMB453, OCUBM and HCC2185).

Moreover, two ILC/ILC-like cell lines (CAMA1 and MDAMB453) had molecular alterations in *PTEN*, both this gene and *PIK3CA* are components of *PI3K/AKT* pathway, one of the most altered pathways in ILC patient tumors ^3,58^.

The final criterion for rational selection is based on deregulated pathways. Often non-recurrent alterations may occur in multiple different components of the same oncogenic pathway resulting in its deregulation. Frequently deregulated oncogenic pathways that are druggable in breast cancer include growth factor and receptor tyrosine kinase (RTK) signaling, cell cycle and TP53 pathway and DNA damage repair (DDR) pathway ^3,58^. Apart from these genomically altered pathways, hormone (ER/PR) signaling is also an important breast cancer pathway that primarily gets deregulated by increase in estrogen hormone levels ^97^. Most of the luminal ILC/ILC-like cell lines showed higher expression of ERpos gene signature and lacked expression of ERneg gene signature indicative of active hormone signaling. In line with these findings, these cell lines also showed higher predicted sensitivity to endocrine therapy (Figure 1C). These findings reveal additional models (such as UACC3133, ZR7530, HCC2218, MDAMB453, WCRC25, and IPH926) to study endocrine response and resistance in human ILC disease. Furthermore, all ILC/ILC-like cell lines had alterations in at least one of key oncogenic breast cancer pathways (Supplementary Figure S12). RTK pathway was altered in all ILC/ILC-like most frequently due to alterations in *ERBB2* and *NF1* gene. Cell cycle pathway was altered in all Lum ILC/ILC-like cell lines due to CCND1 amplifications and CDKN2B mutation. While DDR and PI3K pathways were altered frequently in Lum/HER2 cell lines. With this information, researchers can choose suitable ILC/ILC-like cell lines based on the deregulated pathway on interest. Notably, each of these pathways is druggable using systematic or targeted agents e.g., endocrine therapies to against hormone signaling, HER2 and PI3K inhibitors against RTK signaling ^98,99^, chemotherapy and CDK4/6 inhibitors ^54^ for inhibiting cell cycle and proliferation and PARP inhibitors against tumors with homologous repair defects (HRD) ^100^. Knowledge of these deregulated pathways in ILC/ILC-like cell lines will be critical for developing precision therapies for human-ILC disease.

Finally, in addition to the aforementioned qualitative criterions for choosing ILC/ILC-like cell lines, we also defined quantitative scores for molecular similarity of cell line to ER+ ILC patient tumors to aid in their selection. The molecular similarity of each ILC/ILC-like cell lines to ER+ ILC patient tumors was computed using Pearson correlation of their genome-wide copy number profiles and RNA expression. BCK4 showed the highest copy number and RNA similarity to ILC patient tumors. Most cell lines showed high copy number similarity except for MDAMB468. On the other hand, as expected, all non-luminal (MDAMB468, SKBR3, HCC1187) and few luminal (CAMA1, HCC2185 and OCUMB) showed low or non-significant RNA similarity to ILC patient tumors. Suitability of each cell line to model ER+/luminal ILC disease based on their molecular subtypes and molecular (copy number and RNA) similarity is summarized in Figure 6B.

## Discussion

Despite ILC being acknowledged as a disease with distinct biology that necessitates specialized and precision medicine treatments, the further exploration of its molecular alterations with the goal of discovering new treatments has been hindered due to the scarcity of well-characterized cell line models for studying this disease ^8,9^. Previous large-scale cancer cell line characterization studies such as CCLE 2019 ^11^ and Sanger 2016 ^13^ had limited inclusion of and lacked comprehensive characterization of ILC/ILC-like cell lines.

To address this gap, we generated the ILC Cell Line Encyclopedia (ICLE), providing the first comprehensive molecular characterization of ILC and ILC-like cell lines using state-of-the-art multi-omics comprising optical genome mapping, whole-exome sequencing, RNA sequencing, RPPA, SNP array and DNA methylation array. Our approaches were chosen in line with those used in previous large-scale cancer cell line characterization studies ^11,12^. This allowed us to integrate various ICLE datatypes with corresponding published breast cancer cell line multi-omics datasets from Ghandi et al 2019 (CCLE) ^11^, Marcotte et al 2016 (Neel Lab) ^12^ and Iorio et al 2016 (Sanger) ^13,15^. This allowed us to perform integrative and comparative (ILC vs NST) analyses downstream.

As expected, all the ILC/ILC-like cell lines harbored molecular alterations in *CDH1*/E-cadherin and *CTNNA1*/α-catenin and showed disrupted adherens junctions, a hallmark of ILC phenotype. Moreover, consensus molecular subtyping not only confirmed luminal status of well-established ILC cell lines (MDAMB134VI, MDAMB330, SUM44PE and BCK4) but also revealed additional cell lines that retained luminal molecular features despite the loss of ER-A/PR expression in some cases (IPH926, HCC2185, WCRC25, HCC2218, MDAMB453 and OCUBM) thus expanding the list of luminal models. However, it is important to note that many of these cell lines also showed high HER2/phospho-HER2 expression (several due to HER2/ERBB2 amplifications) and were classified as Lum/HER2 subtype. Notably, despite this feature, most Lum/HER2 ILC/ILC-like cell lines showed high predicted sensitivity to endocrine therapy. Thus, in addition to the well-established ER+/Lum cell lines, these Lum/HER2 cell lines could also serve as promising models to study molecular mechanism of ILC drug resistance to endocrine therapies – key ILC challenge ^1,3,7,8,24,101–103^. Another important question in ILC research is whether patients with ILC show different efficacy to endocrine treatments.

Preliminary data from Big 1-98 trial study (NCT00004205) comparing two endocrine therapies and indicated that patients with ILC show greater benefit with aromatase inhibitors compared to tamoxifen ^104^. It is important to understand whether other types of endocrine treatments could be suitable for ILC treatment and this could be investigated using the expanded array of ER+ Lum or ER+ Lum/HER2 ILC/ILC-like models described in this study.

Moreover, comparison of cell line molecular features to patient tumors revealed that most Lum and Lum/HER2 cell lines showed RNA and copy number similarity to patient tumors. Similarly, ILC/ILC-like cell lines retained molecular alterations in key ILC genes at similar frequency to both primary and metastatic ILC tumors. Notably, ILC/ILC-like cell lines recapitulated the *CDH1* alteration landscape of ILC patient tumors. Like ILC patient tumors, all *CDH1* mutant ILC/ILC-like cell lines had a truncating mutation along with LOH (key mechanism of biallelic inactivation of *CDH1* gene). Interestingly, we also identified novel *CDH1* large-scale structural deletions using OGM in three ILC/ILC-like cell lines as well as exonic deletions in two ILC/ILC-like cell lines, in both cases along with LOH. Notably, all *CDH1* mutations in ILC/ILC-like cell lines were clonal (100% allele frequency), i.e., acquired early in evolution, and seemed unlikely to have acquired in culture. Converse to these similarities, ILC/ILC-like cell lines showed enrichment of molecular alterations in *TP53*, *ERBB2* and several DNA repair genes ( *BRIP1*, *PPM1D* and *DDX5*) which were significantly less frequent in ILC patient tumors.

Recent studies show that structural variations and complex genomic rearrangements are frequently seen in breast cancer genomes ^77,78,105^. However, whole-genome structural variations and complex events such as chromothripsis have been largely uncharacterized in ILC/ILC-like cell lines. Using OGM, we revealed that ILC/ILC-like cell lines showed varying degrees of genomic instability ranging from very complex genomes with frequent structural variations to others (mostly Lum cell lines) with silent genomes with few structural variations. Notably, we identified frequent chromothripsis events in chromosomes 8, 11 and 17 across ILC/ILC-like cell lines. These unique patterns of chromothripsis were breast cancer specific in line with previous reports ^44,77,78^. Interestingly, we also saw enrichment of gain of expression fusions on these same chromosomes implicating chromothripsis events as drivers of these functional fusions ^44,77,78^. Indeed, chromosome 17 contains many important breast cancer genes such as *TP53*, *ERBB2*, *NF1*, *BRCA1*, *MAP2K4*, *PPM1D*, *BRIP1* among others and single catastrophic events like chromothripsis affect large regions (containing various oncogenes and tumor suppressors) providing tumors with opportunities for rapid genome evolution^44,106,107^.

*TFAPAB* gene is an emerging biomarker of lobular disease that is preferentially expressed in both precursor and invasive stages ^89,90^. Importantly, it controls differentiation and proliferation tumor cells and has potential to serve as prognostic biomarker or a targetable vulnerability in ILC disease ^108,109^. For a long time, it was unclear what drives its preferential expression in ILC. Using integrative analysis of DNAm and RNA expression, we revealed epigenetic activation of *TFAP2B* gene in both ILC/ILC-like breast cancer cell lines and ILC patient tumors. This is an important finding that raises questions about whether E-cadherin defects reprogram epigenetic landscape of tumors cells resulting epigenetic activation of proliferation regulators such as *TFAP2B*. Additional studies are warranted to test this hypothesis and further investigate the role of *TFAP2B* in E-cadherin defective breast cancers.

E-cadherin deficiency is an attractive target for development of precision therapies for ILC disease ^92,93,95^. Towards the goal of identifying novel druggable vulnerabilities or dependencies associated with human-ILC disease, we analyzed publicly available RNAi loss of function breast cancer cell line datasets. In line with previous findings, we also observed that ILC/ILC-like cell lines have high dependency genes in cytoskeletal components and PI3K/AKT pathway ^92,93^ further confirming the significance of these pathways as putative vulnerabilities for ILC disease. In addition, we also uncovered novel dependencies in Focal adhesion and various metabolism related pathways related with tyrosine, vitamin B6 and retinoic acid metabolism. Moreover, the top candidate based on druggability score was *PKN3* gene which also showed high expression in ILC vs NST patient tumors (data not shown) and warrant further investigation. Importantly, this is the most comprehensive report on identification of novel druggable vulnerabilities using luminal ILC/ILC-like cell lines and overcomes limitations of previous studies, that due to lack of suitable models, had employed either genetically modified models or basal subtype E-cadherin deficient models to identify druggable vulnerabilities and lacked ILC context.

In summary, we performed comprehensive multi-omic profiling of E-cadherin deficient (ILC/ILC-like) cell lines and highlight their key molecular features including molecular subtypes and genomic alterations, evaluate their similarity to E-cadherin deficient (ILC) breast cancers, catalogue druggable vulnerabilities and finally present a scheme for rational-model-selection. The findings of this study expand the array of suitable cell lines available for modeling human-ILC disease and lay foundation for studying key ILC challenges including development of precision therapies and novel biomarkers as well as for generating novel hypothesis for future ILC research.

## Acknowledgements

We thank Beth Knapick & Jian Chen for their excellent technical support, associated research cores (Bionano Core, UPMC Genome Center, Genome Research Center, Arizona Genetics Core, and MD-Anderson RPPA Core - Grant # 5 P30 CA016672-40), and various funders (FWW, Susan G. Komen, BCRF, Dynami, Magee Womens Research and Education Fund, Shear Family Foundation and NCI R01CA252378, NIH S10OD028483 & others).

## Supplementary Tables Figures

**Supplementary Figure S1.**
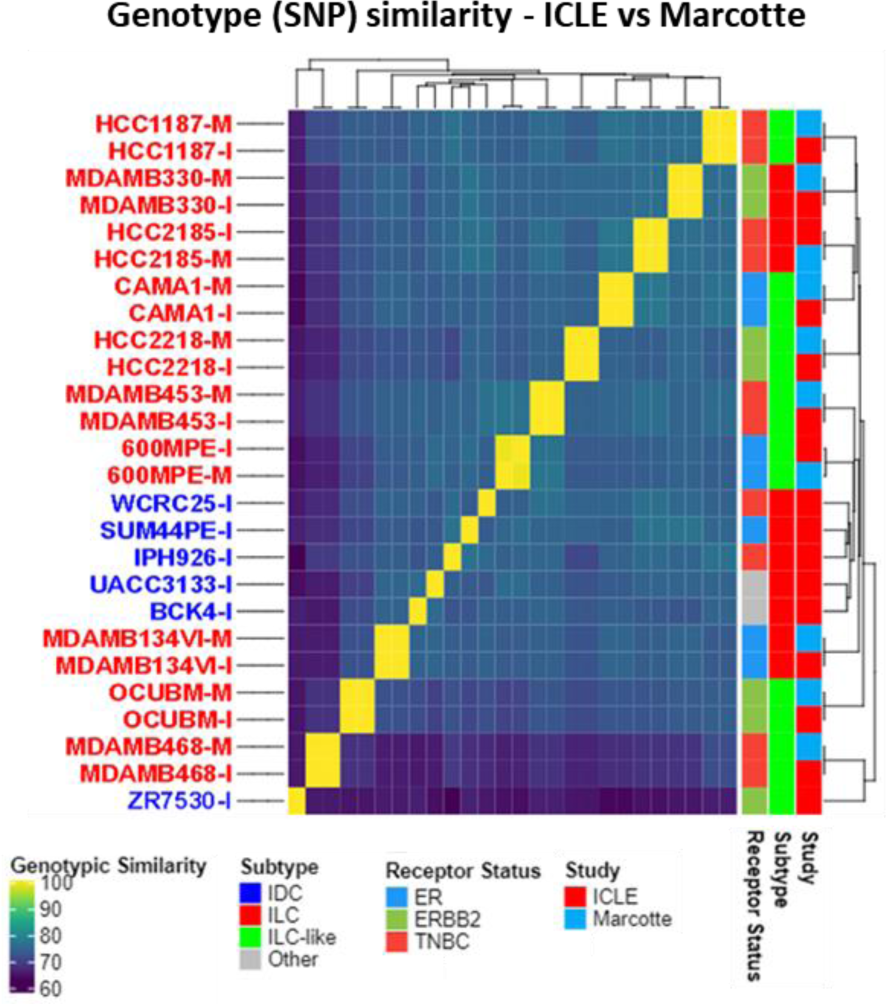
Genotype Similarity between ICLE and Marcotte et al (Neel Lab) Datasets. Five thousand randomly selected SNP calls were compared between all cell lines to assess genotyping similarity (percentage of overlapping SNP calls divided by total SNP calls – 100% means high similarity while 0% means no similarity). Overlapping cell lines are shown in red, while ICLE cell lines with no match in Marcotte dataset are shown in blue.

**Supplementary Figure S2.**
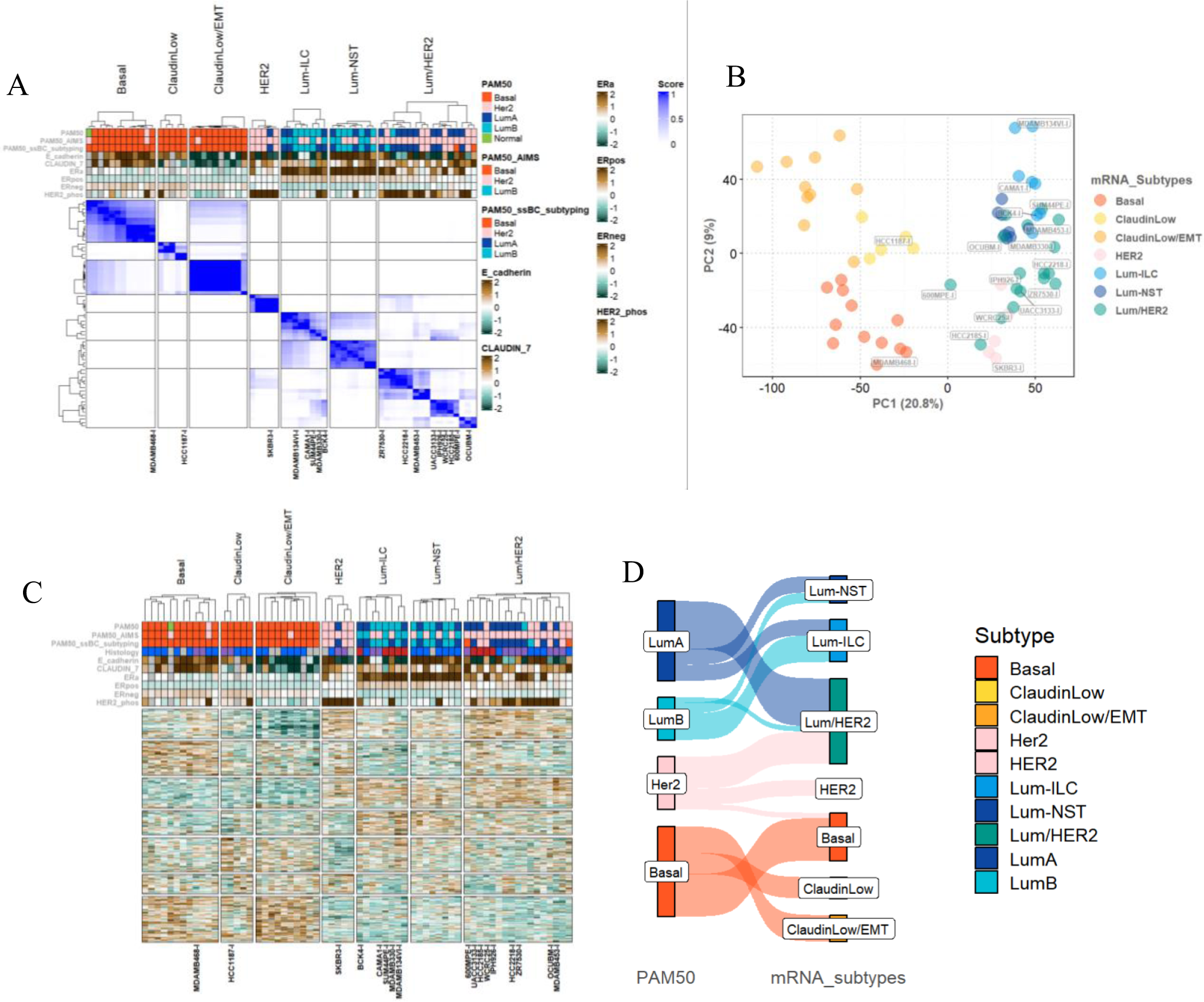
Exploration of RNA Sequencing Dataset and Defining mRNA Subtypes. (A) Consensus clustering was performed on row normalized expression matrix where features represent top 5000 variable features (identified using mean absolute deviation (MAD). K = 9 was used to define refined clusters. Final mRNA subtypes based molecular features of cell lines including E-cadherin, ER, HER2 and Claudin-7 protein levels, ERpos gene signature scores and PAM50 based intrinsic molecular subtypes. Few clusters showing similar molecular features were combined to result in biologically meaningful mRNA subtypes. (B) PCA plot and (C) expression heatmap generated using top variable features show to see transcriptic relationship between distinct mRNA subtypes. (D) Sankey diagram showing relationship between cell line PAM50 calls and mRNA subtypes

**Supplementary Figure S3.**
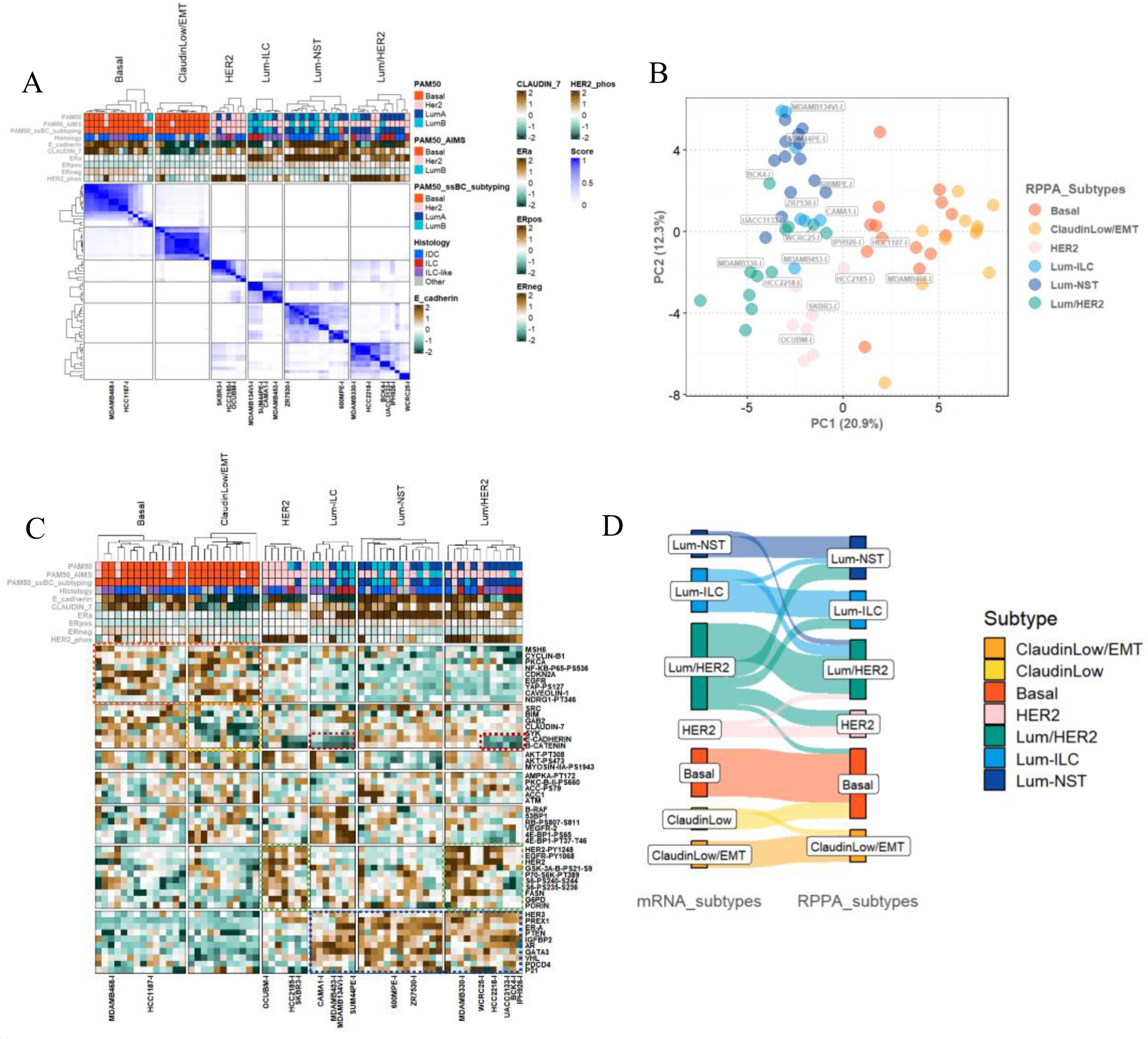
Exploration of RPPA Dataset and Defining RPPA Subtypes. (A) RPPA subtypes defined by consensus clustering of top 50 varibale features defined by MAD as described before. K = 9 was used like before and then based on molecular similarity some clusters were combined to result in 6 clusters. (B) PCA plot and (C) RPPA expression heatmap generated using top variable features show to see proteomic relationship between distinct RPPA subtypes. (D) Sankey diagram showing relationship between cell line mRNA and RPPA subtypes.

**Supplementary Figure S4.**
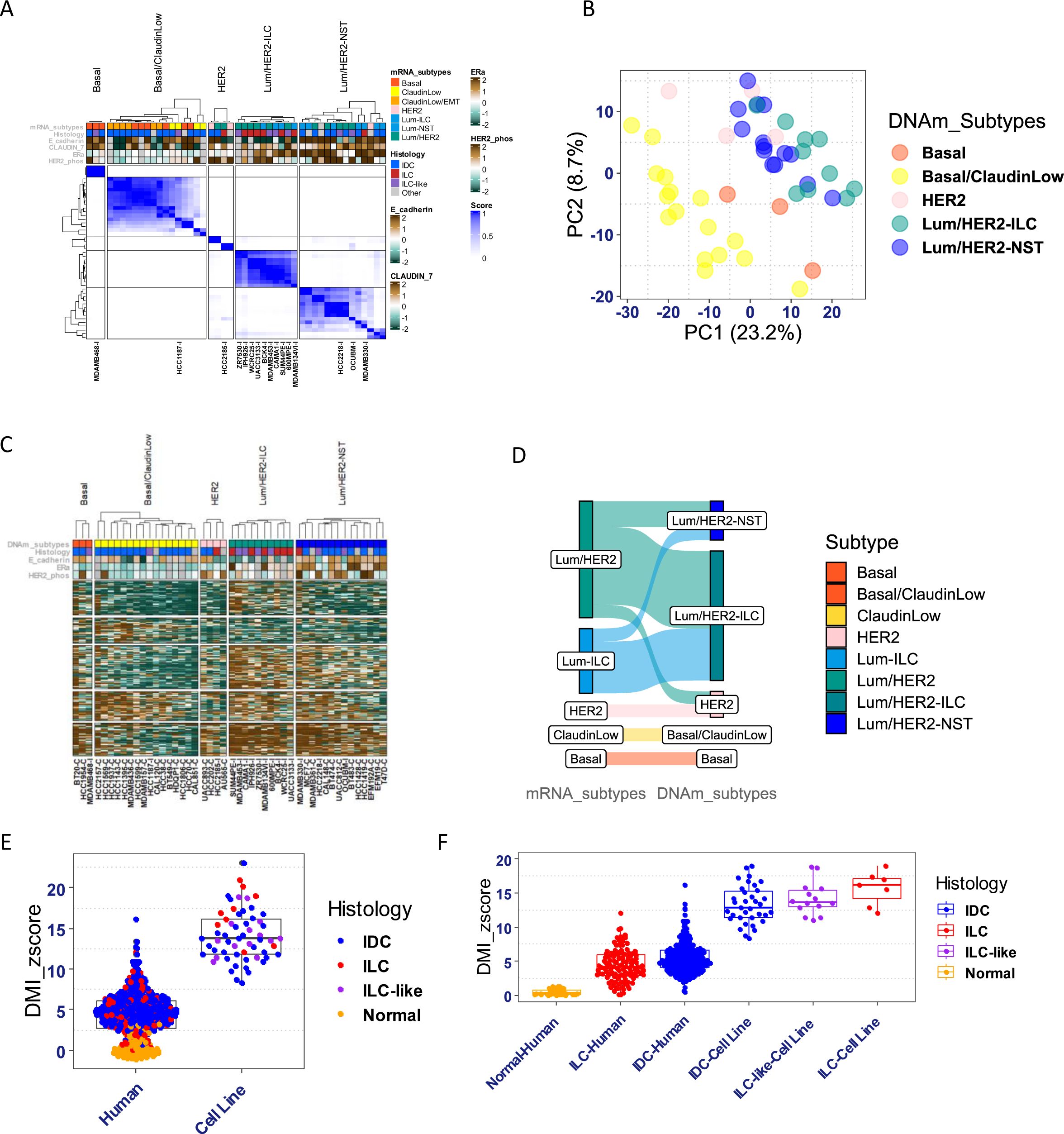
Exploration of DNA Methylation (DNAm) Dataset and Defining DNAm Subtypes. (A) DNAm consensus clustering, top 10% variable feature were selected using variance. DNAm subtypes were defined as desribed for mRNA and RPPA subtypes. (B) PCA plot and (C) DNAm heatmap generated using top variable features show to see DNAm relationship between distinct DNAm subtypes. (D) Sankey diagram showing relationship between cell line mRNA and DNAm subtypes. (E) DNA methylation instability score in BRCA patient tumors (TCGA) vs. BRCA cell lines (ICLE and Sanger) and (F) DNA methylation instability score across different histological subtypes.

**Supplementary Figure S5.**
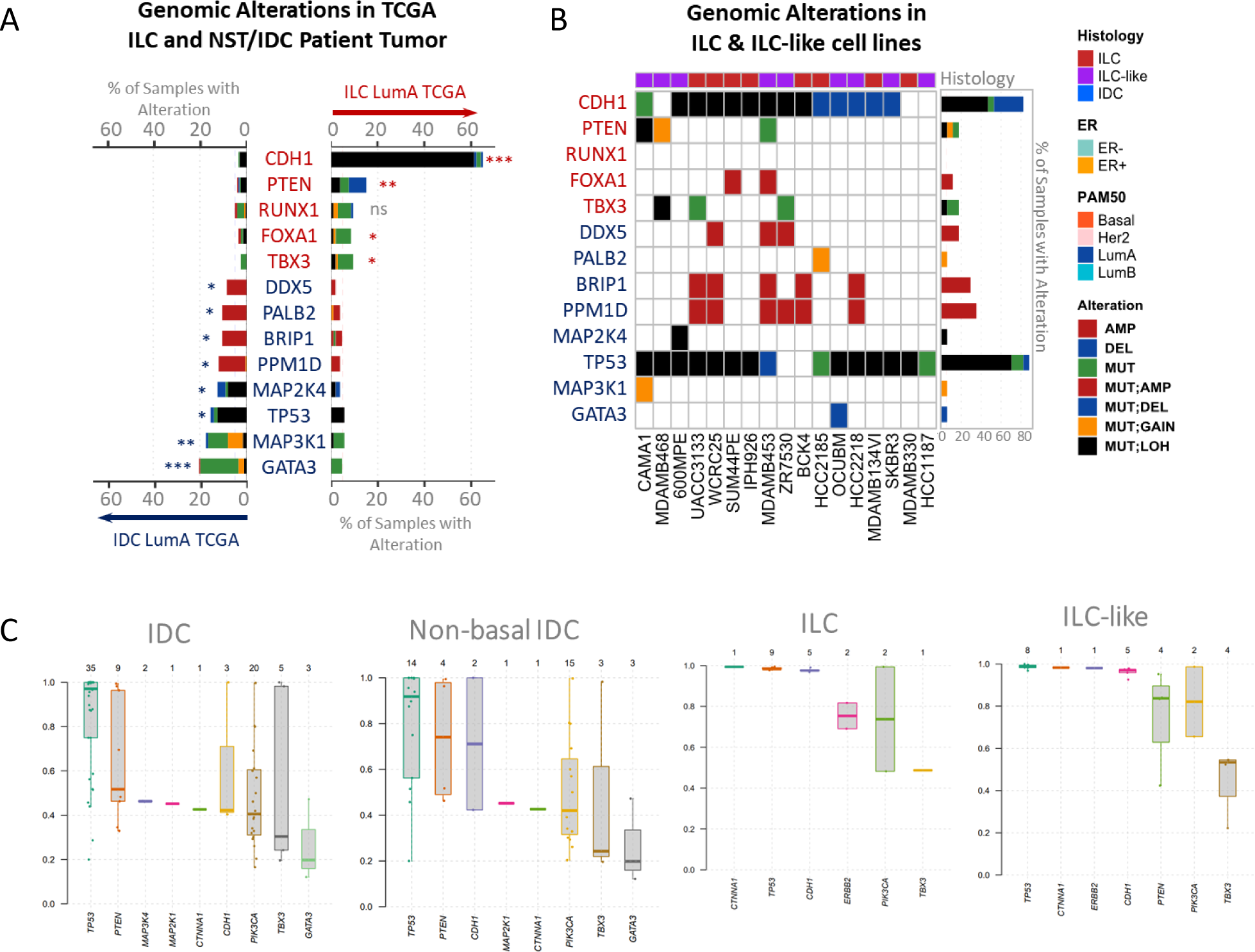
Overview of Key ILC and NST Molecular Alterations. (A) Key Molecular Alterations Enriched in Luminal-A ILC and NST patient tumors in TCGA dataset. (B) Frequency of these alterations in ILC/ILC-like cell lines. (C) Allele frequency of these alterations in all NST/IDC, non-basal NST/IDC, ILC and ILC-like cell lines.

**Supplementary Figure S6.**
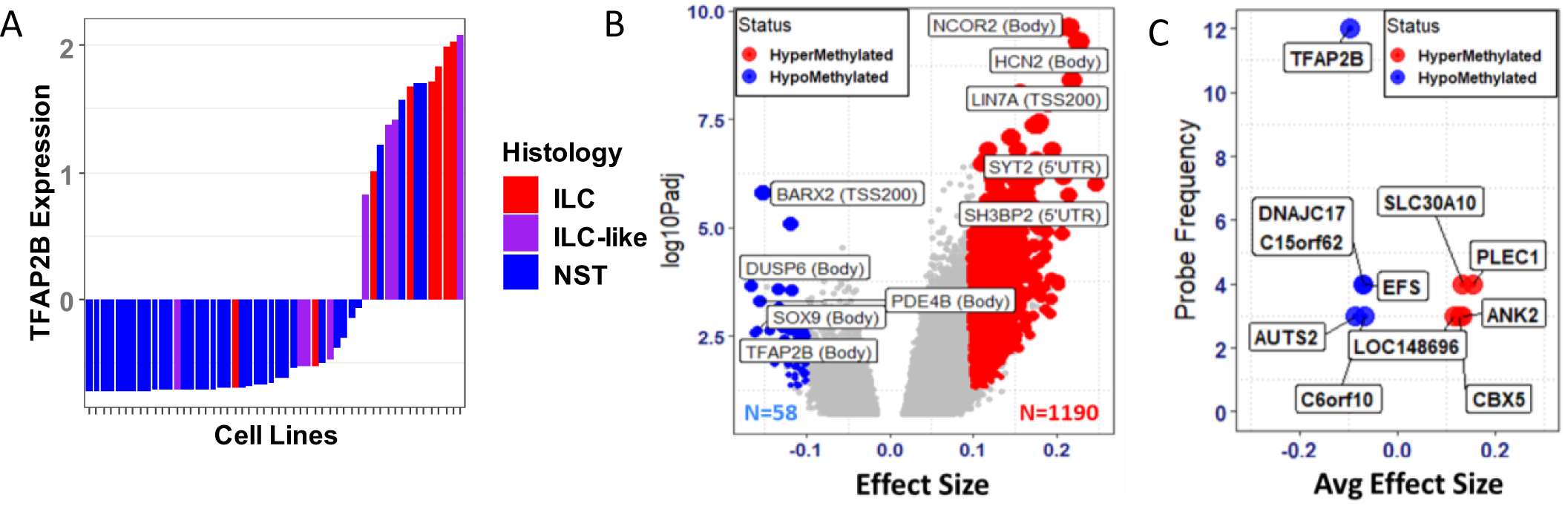
Differential DNA Methylation Analysis of TCGA Luminal-A ILC vs NST Tumors. (A) TFAP2B expression in ILC, ILC-like and NST cell lines showing most ILC/ILC-like genes have high TFAP2B expression. (B) Volcano plot showing differentially methylated probes in TCGA Luminal A ILC vs NST patient tumors annotated by associated gene names (and region). (C) Genes with most frequently methylated probes from panel B. Supplementary Figure S7. p120-catenin, E-cadherin and Hoechst immunofluorescence staining of 17 ILC/ILC-like cell lines.

**Supplementary Figure S7.**
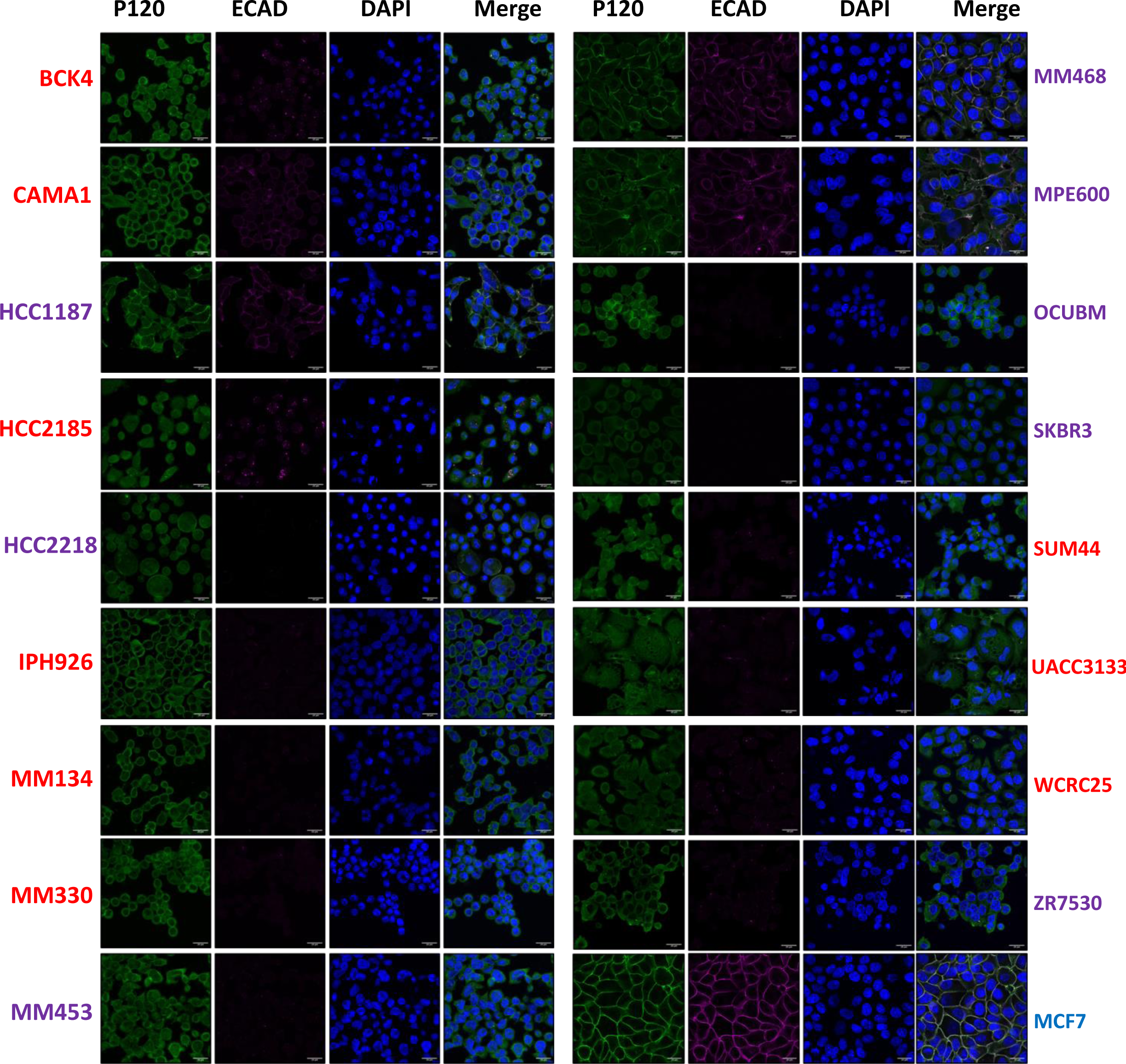
p120-catenin, E-cadherin and Hoechst immunofluorescence staining of 17 ILC/ILC-like cell lines. Immunofluorescence of p120-catenin (1^st^ column), E-cadherin (2^nd^ column), Hoechst nuclear staining (3^rd^ column) and merge of these stainings (4^th^ column) across ILC (names in red), ILC-like (names in purple) and NST (name in blue) cell lines. All ILC/ILC-like cell lines had *CDH1* and *CTNNA1* gene defects resulting in loss of E-cadherin/α- catenin and relocalized of p120-catenin from memberane to cytoplasm. For comparison, we show a CDH1/CTNNA1 WT NST cell line (MCF7) with memberanous staining of both p120 and E-cadherin.

**Supplementary Figure S8.**
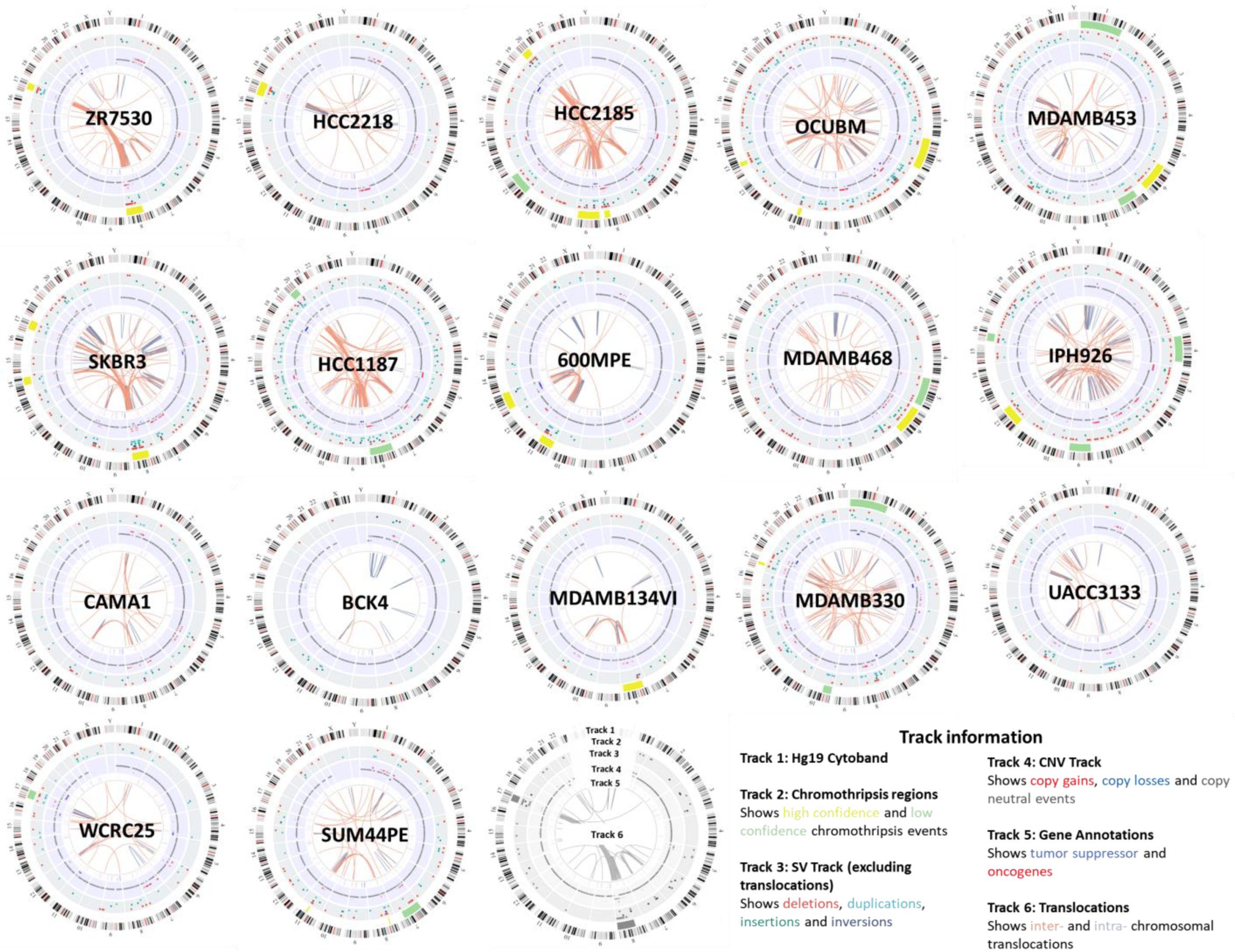
Whole-Genome Landscape of ILC/ILC-like Cell Lines. Circos plots showing OGM identified structural rearrangments, SNP array based copy number, and shatterseek identified chromothripsis events across 17 ILC/ILC-like Cell Lines.

**Supplementary Figure S9.**
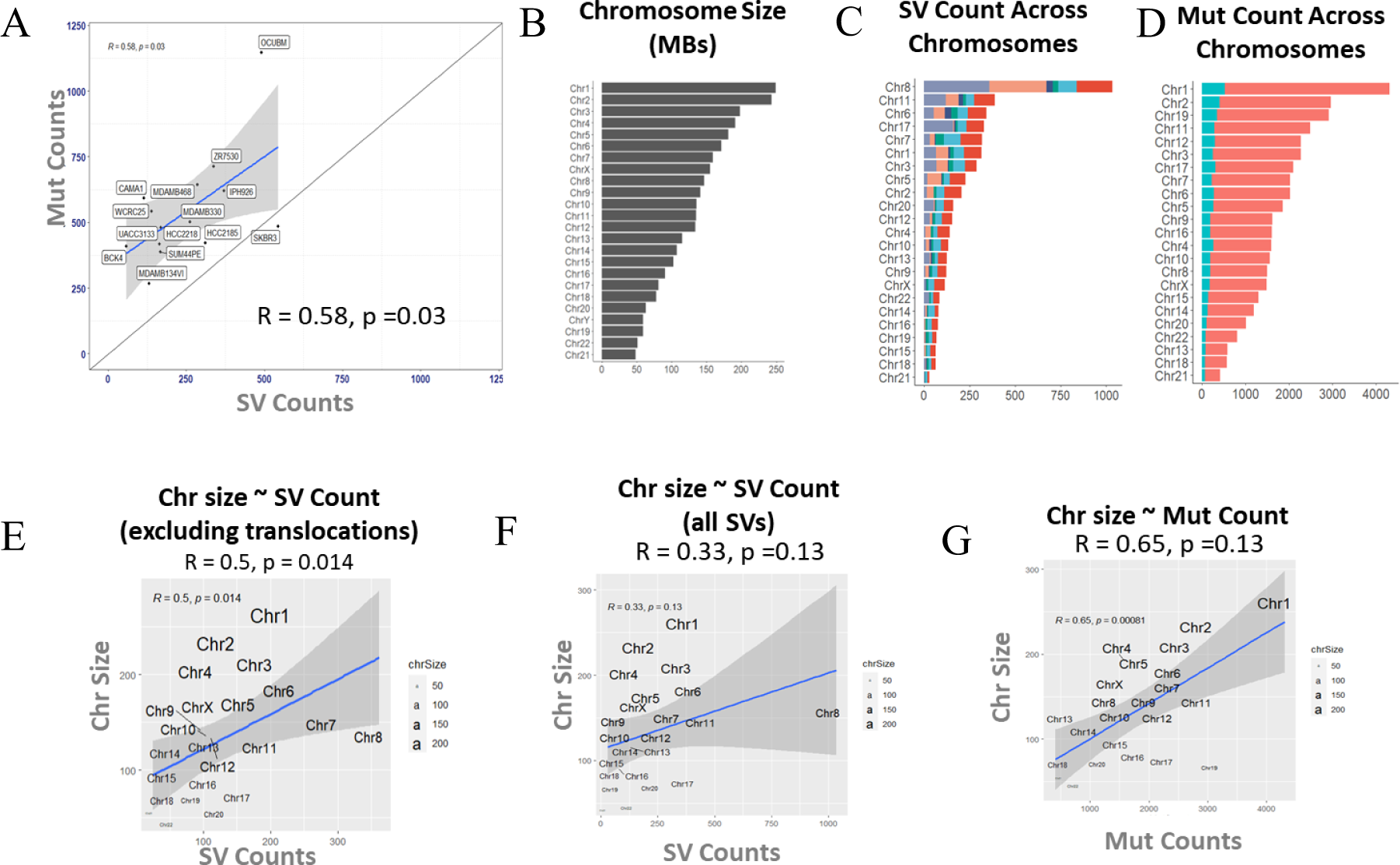
Association Between Various Genomic Metrics of Genomic Instability. (A) Pearson Correlation between SNV/Mutation count and SV count. (B) Barplot showing chromosome size in MBs (megabases). (C) SV count across chromosomes. (D) Mutation count across chromosomes. (E) Pearson correlation between chromosome size and SV count (excluding translocations). (F) Pearson correlation bewtween chromosome size and SV counts. (G) Pearson correlation between chromosome size and mutation count.

**Supplementary Figure S10.**
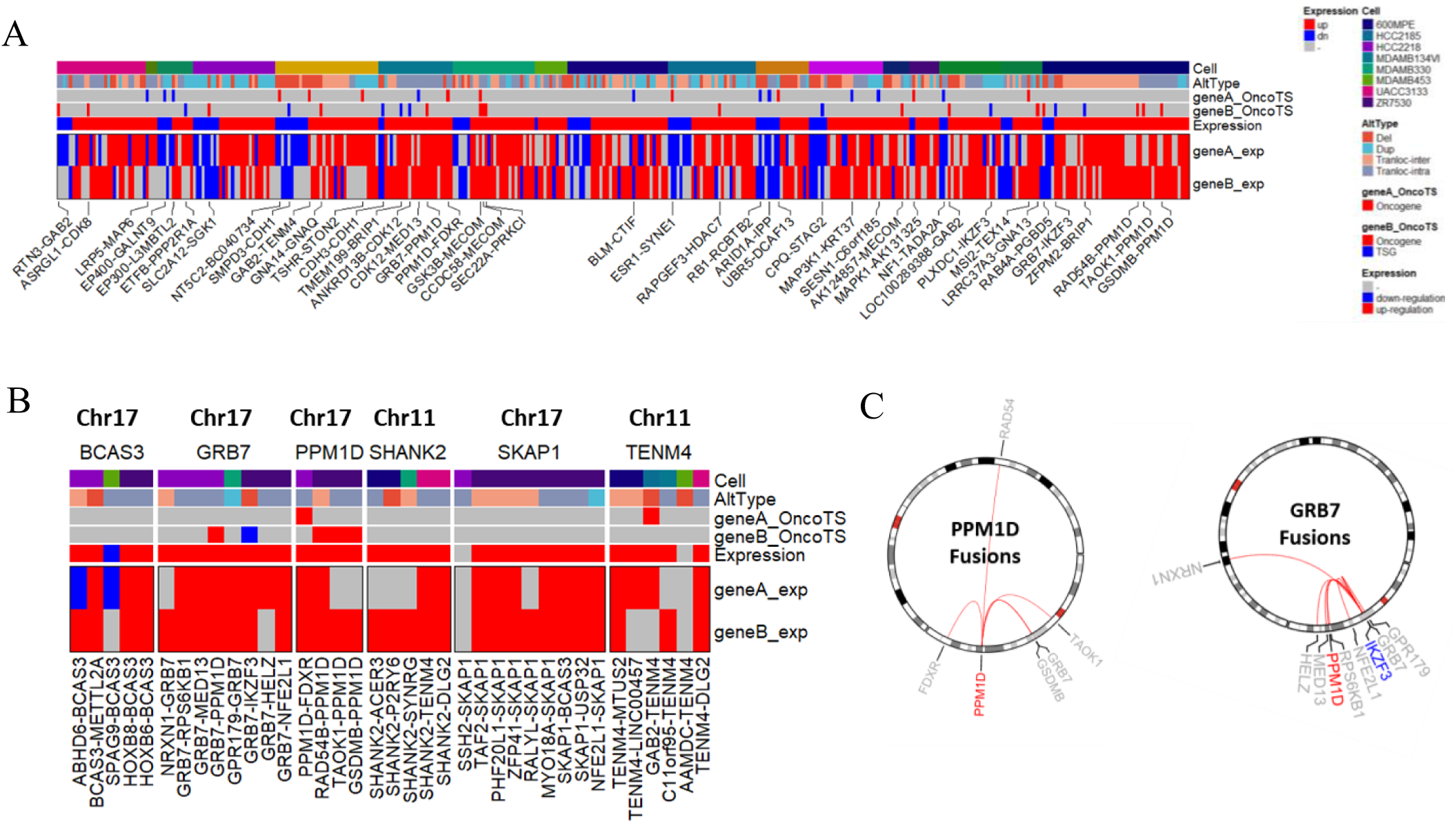
Overview of Functional Fusions. (A) Functional (gain of expression or loss of expression) fusions identified via integrative analysis of OGM and RNAseq datasets across 17 ILC/ILC-like cell lines. Each fusion pair is annotated by cell line, structural alteration type (AltType), fusion gene A and B annotation as oncogene or tumor suppressor (OncoTS) and affect on gene A or gene B expression (B) Recurrently fused genes including BCAS3, GRB7, PPM1D, SHANK2, SKAPI and TENM4. (C) Recurrent fusions of PPM1D and GRB7 genes.

**Supplementary Figure S11.**
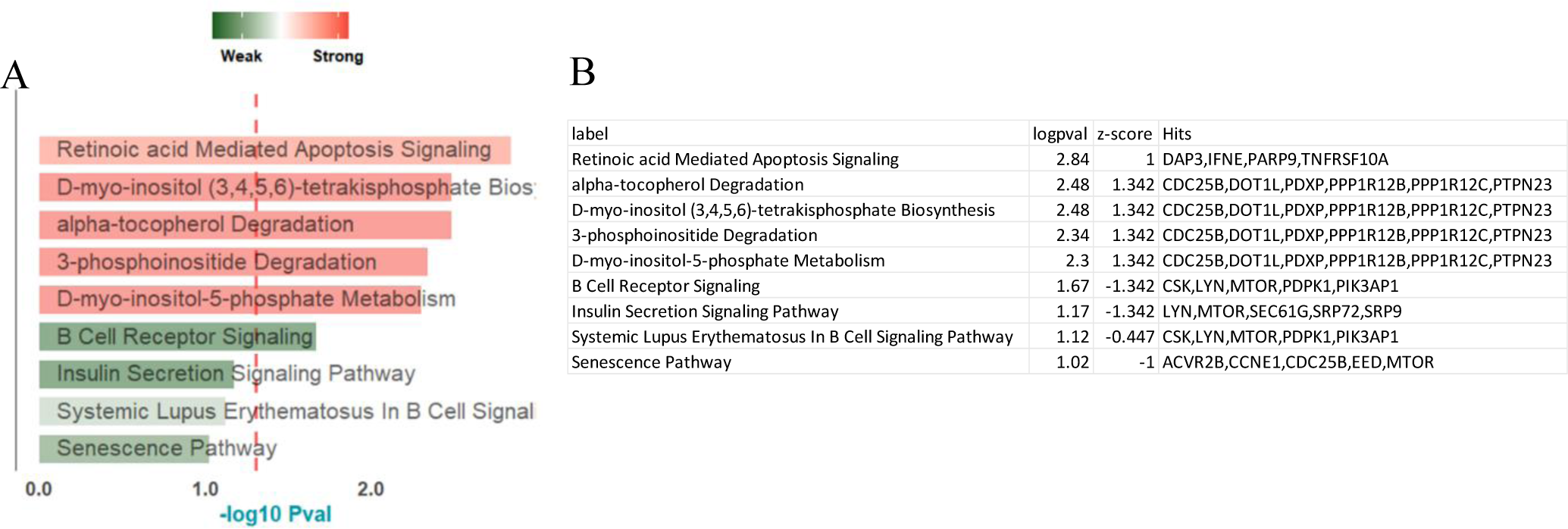
Ingenuity Pathway Analysis (IPA) of Consensus Differential Dependencies. (A) The top strong/high dependency and weak/low dependency pathways identified IPA analysis of consensus differential dependencies. (B) Table showing genes (last column) associated with the pathways shown in A.

**Supplementary Figure S12.**
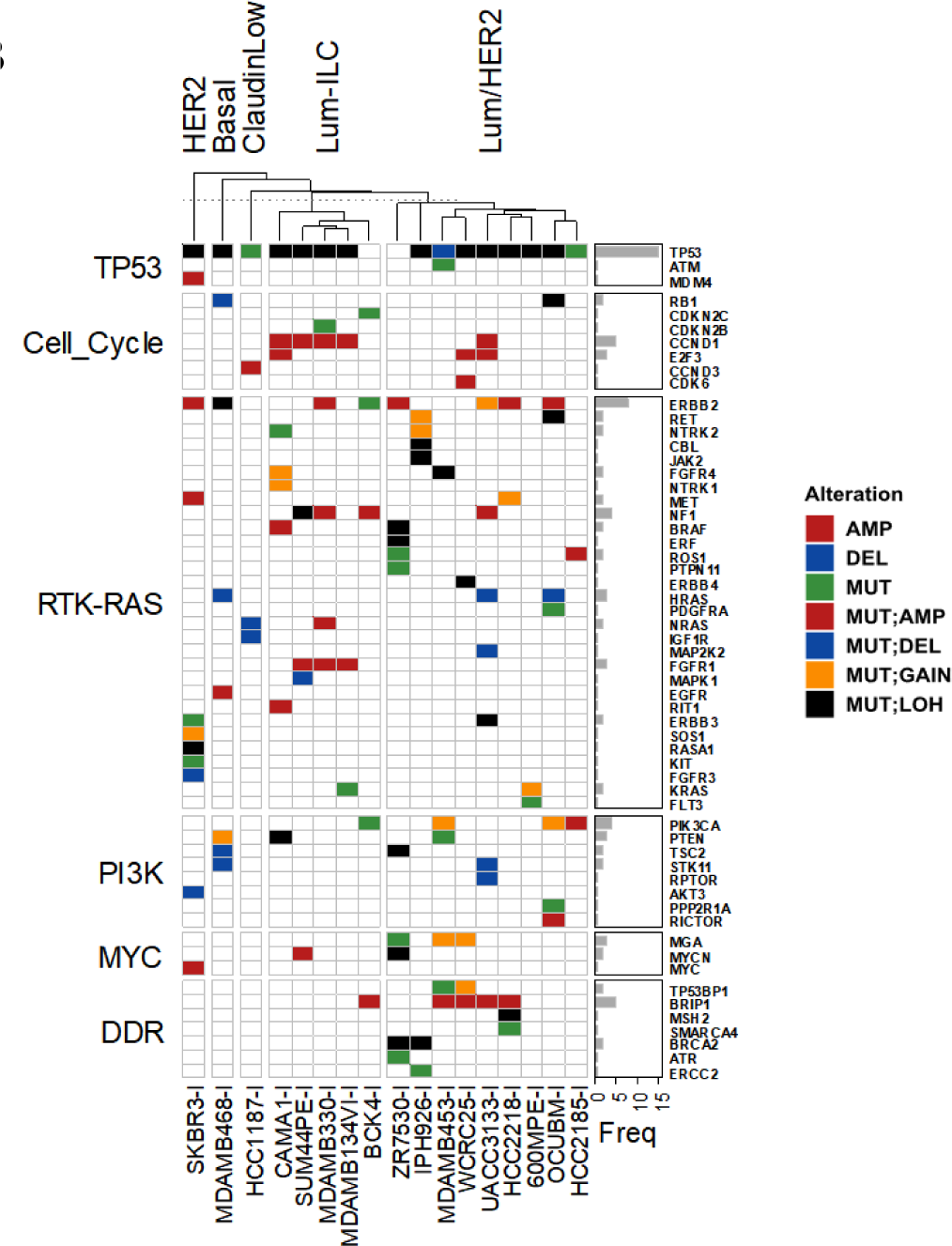
Hormone Signaling Activity and Molecular Alterations in Key Breast Cancer Pathways in ILC/ILC-like cell lines. Molecular alterations in key breast cancer pathways including (Cell Cycle and TP53 pathway, RTK-RAS pathway, PI3K/AKT pathway and DNA damage repair (DDR) pathway genes).

**Supplementary Table S1.** Overview of ICLE Cell line Panel and Their Key Multi-omic Features. Sheet “S1A ICLE Panel Overview” shows clinical charactersitics of the patients from whom the cells were derived, clinical markers, mutational status of ILC driver genes i.e., *CDH1* and CTNNA1, inclusion of these cell lines in previous studies, pubmed report count, key publication/reference, cellosaurus, availability and media conditions. Sheet S1B “ICLE Multi-omic Features” shows receptor subtypes, molecular subtypes, molecular alterations, genomic instability metrics and molecular resemblence to ILC patient tumors. Receptor subtypes are based on ER, PR and HER2 protein (RPPA) levels., metrics of genomic instability, molecular resemblence to ILC patient tumors and cell line classification. Molecular Subtypes include PAM50 based subtypes and consensus clustering based multi-omic subtypes (mRNA subtypes, RPPA subtypes and DNAm subtypes). Molecular Alterations are shown for key ILC drivers genes (CDH1 and CTNNA1) and other ESCAT actionable genes. Genome-wide metrics include fraction of genome altered (FGA), tumor mutation burden (TMB), structural variation count (SV), and chromothripsis events. Molecular resemblence is based on correlation of all ICLE cell line copy number and mRNA expression profiles with those of ILC patient tumors from TCGA cohort. Finally, the ICLE cell line classification is based on consideration of molecular subtypes and molecular resmeblence to ILC patient tumors.

| Cell Line | All PubMed Reports (June 2023) | PubMed Reports focused on ILC (June 2023) | Type | Receptor Subtype | Protein Levels |  |  |  | Molecular Subtypes |  |  |  | ILC Drivers |  | Genomic Instability Metrics |  |  |  |  |
| --- | --- | --- | --- | --- | --- | --- | --- | --- | --- | --- | --- | --- | --- | --- | --- | --- | --- | --- | --- |
|  |  |  |  |  | ER | PR | HER2 | HER2_Py1248 | PAN50 | mRNA Subtype | RPPA Subtypes | DNAm Subtypes | CDH1 | CTNNA1 | FGA | TMB | SV | DMI | Chromothripsis |
| BCK4 | 7 | 6 | ILC | ER+ | High | High | Mid | High | LumA | Lum | Lum/HER2 | Lum/HER2 | MUT1,LOH |  | 0.14 | 5.46 | 111 | 0.75 |  |
| HCC2185 | 0 | 0 | ILC | TN |  | Low | Mid | Mid | LumA | Lum/HER2 | HER2 | Lum/HER2 | DEL |  | 0.25 | 5.20 | 348 | 0.54 |  |
| IPH926 | 7 | 6 | ILC | TN | Mid | Low | Mid | Mid | LumA | Lum/HER2 | Lum/HER2 | Lum/HER2 | MUT1,LOH |  | 0.37 | 7.82 | 370 | 0.71 |  |
| MDAMB134VI | 51 | 11 | ILC | ER+ | High | Low | Low | Mid | LumB | Lum | Lum | Lum/HER2 | EXON6,LOH |  | 0.18 | 3.66 | 131 | 0.91 | chr8 |
| MDAMB330 | 20 | 2 | ILC | ER/HER2+ | High | Low | High | High | LumB | Lum | Lum/HER2 | Lum/HER2 |  | MUT | 0.27 | 6.92 | 262 | 0.77 |  |
| SUM44PE | 26 | 10 | ILC | ER+ | High | Low | Low | Low | LumB | Lum | Lum | Lum/HER2 | MUT1,LOH |  | 0.28 | 4.56 | 167 | 0.88 | chr8, chr11 |
| UACC3133 | 1 | 0 | ILC | ER/HER2+ | High | Low | Mid | High | LumA | Lum/HER2 | Lum/HER2 | Lum/HER2 | MUT1,LOH |  | 0.33 | 5.14 | 228 | 0.58 |  |
| WCRC25 | 0 | 0 | ILC | TN | Low |  | Mid | Low | Her2 | Lum/HER2 | Lum | Lum/HER2 | MUT1,LOH |  | 0.14 | 7.02 | 139 | 0.69 |  |
| 600MPE | 7 | 0 | ILC-like | ER+ | High | Low | Mid |  | LumA | Lum/HER2 | Lum | Lum/HER2 | MUT1,LOH |  | 0.18 | 3.58 | 254 | 0.52 | chr11, chr13 |
| CAMA1 | 69 | 0 | ILC-like | ER+ | High | High | Mid |  | LumB | Lum | Lum | Lum/HER2 | MUT |  | 0.26 | 8.20 | 158 | 0.70 |  |
| HCC1187 | 12 | 0 | ILC-like | TN |  | Low | Mid |  | Basal | ClaudinLow | Basal | ClaudinLow/EMT |  | MUT | 0.21 | 4.38 | 256 | 0.52 | chr3 |
| HCC2218 | 6 | 0 | ILC-like | HER2+ |  | Low | High | High | LumA | Lum/HER2 | Lum/HER2 | Lum/HER2 | DEL |  | 0.24 | 6.98 | 168 | 0.62 | chr17 |
| MDAMB453 | 676 | 3 | ILC-like | HER2+ | Mid | Low | High | High | LumA | Lum/HER2 | Lum | Lum/HER2 | MUT1,LOH |  | 0.23 | ### | 270 | 0.82 | chr6 |
| MDAMB468 | 2200 | 4 | ILC-like | TN | Low | Low | Low | Mid | Basal | Basal | Basal | Basal |  | EXON4,5,MUT | 0.23 | 7.30 | 335 | 0.64 |  |
| OCUB-M | 5 | 0 | ILC-like | HER2+ | Low | Low | High | High | Her2 | Lum/HER2 | HER2 | Lum/HER2 | EXON2,LOH |  | 0.41 | ### | 551 | 0.58 | chr10 |
| SK-BR-3 | 2925 | 11 | ILC-like | HER2+ | Mid | Low | High | High | LumA | HER2 | HER2 |  | DEL |  | 0.21 | 6.64 | 664 |  | chr17 |
| ZR7530 | 110 | 0 | ILC-like | ER/HER2+ | High | Mid | Mid | High | LumA | Lum/HER2 | Lum | Lum/HER2 | MUT1,LOH |  | 0.21 | 8.06 | 392 | 0.60 | chr17 |

**Supplementary Table S2.** Overview of the Generated ICLE Datasets. Columns show data types, technology used to generate the data and data levels for all 17 ILC/ILC-like cell lines.

| Data Types | Technology | Data Levels | MDA-MB-134-VI | SUM44PE | MDA-MB-330 | IPH926 | WCRC25 | HCC2185 | BCK4 | UACC-3133 | HCC1187 | MPE600 | MDA-MB-453 | HCC2218 | ZR-75-30 | MDA-MB-468 | SK-BR-3 | CAMA-1 | OCUB-M |
| --- | --- | --- | --- | --- | --- | --- | --- | --- | --- | --- | --- | --- | --- | --- | --- | --- | --- | --- | --- |
| Structural Variations (SV) | Bionano Optical Mapping | L1: Molecules,<br>L2: Variant Report | ✓ | ✓ | ✓ | ✓ | ✓ | ✓ | ✓ | ✓ | ✓ | ✓ | ✓ | ✓ | ✓ | ✓ | ✓ | ✓ | ✓ |
| Single Nucleotide Variations (SNV) | Whole Exome Sequencing<br>(xGen Exome Research Panel v1.0) | L1: Fastq, L2: BAM,<br>L3: VCF, L4: MAF | ✓ | ✓ | ✓ | ✓ | ✓ | ✓ | ✓ | ✓ | ✓ | ✓ | ✓ | ✓ | ✓ | ✓ | ✓ | ✓ | ✓ |
| Copy Number Variations (CNV) | Illumina CytoSNP 850K v1.2 | L1: Idat, L2: logRR + BAF,<br>L3: Seg file, L4: GISTIC | ✓ | ✓ | ✓ | ✓ | ✓ | ✓ | ✓ | ✓ | ✓ | ✓ | ✓ | ✓ | ✓ | ✓ | ✓ | ✗ | ✓ |
| Reverse Phase Protein Array (RPPA) | MD-Anderson RPPA | L1: Raw Intensities,<br>L2: Normalized Intensities | ✓ | ✓ | ✓ | ✓ | ✓ | ✓ | ✓ | ✓ | ✓ | ✓ | ✓ | ✓ | ✓ | ✓ | ✓ | ✓ | ✓ |
| DNA Methylation Array (DNAm) | Infinium MethylationEPIC | L1: Idat, L2: B-value | ✓ | ✓ | ✓ | ✓ | ✓ | ✓ | ✓ | ✓ | ✓ | ✓ | ✓ | ✓ | ✓ | ✓ | ✓ | ✗ | ✓ |
| RNA Sequencing (RNAseq) | Poly-A RNA Sequencing<br>(KAPA RNA hyperPrep kit) | L1: Fastq, L2: BAM/quant,<br>L3: Counts | ✓ | ✓ | ✓ | ✓ | ✓ | ✓ | ✓ | ✓ | ✓ | ✓ | ✓ | ✓ | ✓ | ✓ | ✓ | ✓ | ✓ |

**Supplementary Table S3.** Overview of Public Datasets Used for Intergration. Columns show ICLE datasets and corresponding public datasets used for intergration across each datatype.

| ICLE Data Sets<br>(Technology) | Public Data Sets Used for Integration<br>(Technology) |
| --- | --- |
| SV<br>(Bionano Optical Mapping) | - |
| SNV<br>(xGen Exome Research Panel v1.0) | Ghandi et al 2019 (CCLE)<br>(WES, WGS, RNAseq) |
| CNV<br>(Illumina CytoSNP 850K v1.2) | Marcotte et al 2016 (Neel Lab)<br>(Illumina Omni Quad) |
| RPPA<br>(MD-Anderson RPPA) | Marcotte et al 2016 (Neel Lab)<br>(MD-Anderson RPPA) |
| DNAm Array<br>(Infinium MethylationEPIC) | Iorio et al 2016 (Sanger)<br>(Infinium Methylation 450K) |
| RNAseq<br>(KAPA RNA hyperPrep kit) | Ghandi et al 2019 (CCLE)<br>(Poly-A RNAseq) |

